# Competition for water and species coexistence in phenologically structured annual plant communities

**DOI:** 10.1101/2021.10.15.464433

**Authors:** Jacob I. Levine, Jonathan M. Levine, Theo Gibbs, Stephen W. Pacala

**Affiliations:** Department of Ecology and Evolutionary Biology, Princeton University, Princeton NJ, USA; Lewis-Sigler Institute for Integrative Genomics, Princeton University, Princeton NJ, USA

## Abstract

Both competition for water and phenological variation are important determinants of plant community structure, but ecologists lack a synthetic theory for how they affect coexistence outcomes. We developed an analytically tractable model of water competition for Mediterranean annual communities and demonstrate that variation in phenology alone can maintain high diversity in spatially homogenous assemblages of water-limited plants. We modeled a system where all water arrives early in the season and species vary in their ability to grow under drying conditions. As a consequence, species differ in growing season length, and compete by shortening the growing season of their competitors. This model replicates and offers mechanistic explanations for qualitative patterns observed in prior empirical studies of how phenology influences coexistence among Mediterranean annuals. Additionally, we found that a decreasing, concave-up tradeoff between growth rate and access to water can theoretically maintain infinite diversity under simple but realistic assumptions. High diversity is possible because: 1) later plants escape competition after their earlier-season competitors have gone to seed and 2) early-season species are more than compensated for their shortened growing season by a growth-rate advantage. Together, these mechanisms provide an explanation for how annual plant species might coexist when competing only for water.

## Introduction

Competition for water is an important driver of plant community structure globally (Rosenzweig 1968; Fowler 1986; Goldberg & Novoplansky 2009; Craine & Dybzinski 2013; Schwinning & Kelly 2013), yet ecologists lack synthetic theory for how species coexist on limited water resources (Craine & Dybzinski 2013). In place of such theory, community ecologists often use generic consumer resource models with an abiotic “resource” to understand and predict the outcome of water competition among plants (Seabloom *et al*. 2003). However, the use of these models might miss critical aspects of plant competition that uniquely emerge when species compete for water. Indeed, past empirical studies suggest that while consumer-resource models adequately predict the coexistence and competitive dynamics of plants competing for soil nutrients, unexpected dynamics can emerge when plants compete for water (Farrior *et al*. 2013). For example, Farrior et al. (2013) predict that additional nitrogen should decrease a plant’s allocation to roots, but that additions of water should increase root allocation – a dynamic that would be missed in a conventional consumer-resource modeling framework. These predictions were then verified experimentally (Farrior *et al*. 2013). Properly understanding coexistence when plants compete for water might require integrating over the temporally variable physical factors that affect water availability; the physiological relationships between water availability and plant growth and mortality; and demography, the net result of variable plant growth and death rates over time (Daly *et al*. 2004; Rodriguez-Iturbe & Porporato 2007). However, when integrated across multiple competing species, these processes result in intractable, highly nonlinear dynamics.

Recent advances in ecohydrology (Box 1) simplify the problem considerably, and thereby facilitate the development of tractable models of the coexistence of plant species competing for water. Specifically, the equations governing stomatal behavior (Ball *et al*. 1987; Sperry *et al*. 2016; Wolf *et al*. 2016) show that stomatal aperture, and therefore biomass growth, is virtually independent of soil water until soil becomes drier than a species-specific threshold. At this point, a plant’s biomass growth rate and soil water depletion feedback on one another, rendering models of biomass growth mathematically intractable. This remains true until biomass growth falls to zero at a species-specific threshold minimum water availability. What ultimately makes this dynamic tractable is the recent discovery that the period of water limited growth is short enough to be ignored for most competition problems. Specifically, when limited by water plants run the risk of embolism, which if severe, greatly reduces water transport and depresses xylem water potentials. To avoid these costs, plants tend to shut stomates well before substantial embolism risk occurs (Farquhar & Sharkey 1982; Tyree & Sperry 1989; Sala et al. 2012; Anderegg et al. 2016; Wolf et al. 2016), effectively truncating the period of water limited growth. As a result, the relationship between biomass growth rate and soil water availability can be safely approximated as a simple step function – plants grow uninhibited above some species-specific threshold water availability and stop growing below that threshold, a value we call a plant species’ “shutoff point” (Box 1; Appendix 1.1).

### Box 1 Ecophysiology relevant to the modeled dependence of biomass growth on soil water

Photosynthesis requires a continuous supply of water as controlled by a plant’s stomatal conductance, 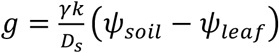, where *γ* is the ratio of root area to leaf area, *k* is the plant’s xylem conductivity and *D_s_* is a measure of air moisture moisture content (vapor pressure deficit). For stomatal conductance g, and therefore photosynthesis, to remain positive there must be a positive gradient between soil water potential, *ψ_soil_* and a plant’s leaf water potential, *ψ_leaf_* (Farquhar & Sharkey 1982). Therefore, as soil water potential drops throughout the season due to loss of soil moisture via evapotranspiration, a plant’s leaf water potential, *ψ_leaf_*, must also drop to maintain a positive gradient. However, *ψ_leaf_* can only reduce to a certain value before embolism risk becomes high. To protect against embolism, plants have an internal shutoff mechanism, *β*(*ψ_leaf_*), which closes stomates, reducing stomatal conductance in order to keep *ψ_leaf_* at a safe level, *ψ*\* (Wolf, Anderegg, and Pacala, 2016). The value of *ψ*\* is typically buffered against values for which embolism is a severe risk and is primarily determined by a plant’s physiological characteristics (Daly *et al*. 2004; Manzoni *et al*. 2011). Above *ψ*\*, the gradient between leaf and soil water potential is large enough that photosynthesis is not water limited — meaning the plant accumulates carbon at its maximum rate, *a_max_*, which is determined by light availability and the plant’s internal carbon concentration (Farquhar and Sharkey 1982; Leuning 1995). Once *ψ_leaf_* = *ψ*\*, *ψ_leaf_* remains fixed, photosynthesis becomes water limited, and carbon accumulation, *α*, drops until the soil water potential also equals *ψ*\* at which point the plant stops growing (Fig. S1).

In the period between when *ψ_leaf_* first reaches *ψ*\* and when the plant shuts down, the dependence of a plant’s growth rate of water content causes demographic models of waterlimited plants to be intractable. However, several considerations lead to a simple approximation. First, a plant’s carbon accumulation rate is a function of stomatal conductance which is itself a function of both the plant’s carbon accumulation rate and *β*(*ψ_leaf_*) (Farquhar and Sharkey 1982; Leuning 1995). The dependence of the carbon accumulation rate on the carbon accumulation rate itself creates a positive feedback such that in the period during which *ψ_leaf_* = *ψ*\* the rate of decrease in a plant’s carbon accumulation is both rapid and accelerating. Second, when plants transpire, they remove water from the soil volumetrically. Volumetric water content is related nonlinearly to soil water potential in a manner such that a given volumetric removal results in a greater reduction of soil water potential in dry conditions than in wet conditions. Additionally, a plant’s evapotranspiration rate increases as a function of plant size. Since plants are largest and soil is driest at the end of the season, the period of declining growth rate is negligible in terms of its contribution to total growth. Detailed calculations supporting this result are provided in Appendix 1.3. Having established that the declining growth rate in this period has a negligible effect on total within-season plant growth, we can safely approximate a plant’s carbon accumulation rate as a step function such that *a* = *a_m_* when *ψ_soil_* > *ψ*\*, and *a* = 0 when *ψ_soil_* ≤ *ψ*\*.

As individual plants grow, they withdraw and transpire water from a common pool at a rate proportional to their leaf area, stomatal conductance and vapor pressure deficit. This relationship is given by

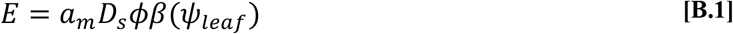

where *E* is the rate of evapotranspiration per unit leaf area, *a_m_* is the maximum photosynthetic rate, internal carbon concentration and the leaf respiration cost, *D_s_* is the vapor pressure deficit, *β*(*ψ_leaf_*) is the shutoff operator, and *ϕ* is a collection of physiological parameters which in addition to *a_m_* determine stomatal conductance (see Appendix 1 for a full description of parameters). Since we are primarily interested in parsing the effects of water-limitation induced variability in phenology, we assume that the constants in Equation B.1 are invariable in time and among species. Thereby, species operate at their maximum photosynthetic rate until 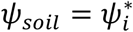, at which point they stop growing and convert to seed. This facet of plant growth also implies that the evapotranspiration rate per unit leaf area remains constant throughout the growing season.

Here we show how the phenological structuring of plant communities that emerges when water is seasonally pulsed and species differ in their shutoff points enables the coexistence of plant competitors. Specifically, species with costly-to-produce thick-walled xylem and high allocation to roots grow longer under drier conditions than species with opposing traits, as these features help plants avoid embolism. Because roots and thick-walled xylem are made at the expense of productive leaves, this creates a tradeoff between biomass growth rate and the minimum soil water availability at which a species can continue to grow (Enquist, Brian *et al*. 1999; Hacke *et al*. 2001; Rosner 2017). This tradeoff then sorts species in time. When the soil is wet, the fastest growing species builds biomass, and hence future fecundity, more rapidly than any other species. But it then stops growing before all others as the soil dries, leaving a period of growth for species able to transpire (and continue to grow) at low water potentials (Appendix 4.2). In sum, interspecific variation in tolerance to low water availability, combined with temporal variation in water supply means that successively smaller subsets of species are actively growing as the season progresses and water is depleted.

Although exactly how this phenological structuring maintains plant species diversity is only beginning to be explored in mechanistic models, we have good reason to believe that phenological variation, water or otherwise, plays a significant role in plant coexistence. First, we commonly observe large phenological diversity in plant communities throughout the world (Rathcke & Lacey 1985; Craine *et al*. 2012). Moreover, phenological differences between species determine the outcome of interactions between invasive and resident plant species (Fridley 2012; Godoy & Levine 2014) and the overyielding potential in diverse communities (Hooper & Dukes 2010). Patterns such as these have motivated past verbal models of how phenological variation influences ecological dynamics, invoking concepts such as phenological niches and niche preemption (Wolkovich & Cleland 2011). Since then, studies have combined empirical work with coexistence theory (Chesson 2000) to explore how phenological differences relate to the niche differences that stabilize coexistence and the competitive imbalances that drive the exclusion of inferior competitors (Godoy & Levine 2014; Alexander & Levine 2019). However, this phenomenological modeling approach relies on high-level abstractions of the competitive process, and does not identify the source of the interactions enabling coexistence. Thus, a detailed, physiologically grounded model might clarify the processes by which phenological structuring in water limited communities influences competitive outcomes.

In this paper we develop theory informed by ecohydrology to answer the following questions: 1) Can tradeoffs associated with water competition explain the high diversity observed in phenologically structured plant communities in nature? 2) Under what conditions do these mechanisms foster coexistence vs. competitive exclusion?

We examine these questions by modeling a system with relatively simple water dynamics, demography, and phenological structure: Mediterranean annual plant assemblages. In these assemblages, a period of winter rainfall and abundant water is followed by a warmer dry period of water limitation, after which, plants successively die. These simplifying features of the hydrology are important because complexity in the inputs and losses of water, as characterize most other systems, makes modeling competition for water particularly challenging (Daly *et al*. 2004; Rodriguez-Iturbe & Porporato 2007). For example, on the input side, precipitation comes in the form of storm events, which vary stochastically in their intensity, frequency and duration (Fernandez-Iliescas & Rodriguez-Iturbe 2003; Rodriguez-Iturbe & Porporato 2007). Systems dependent on stochastic, pulsed inputs interspersed with periods of water limitation make challenging subjects for modeling competition.

By contrast, in Mediterranean annual systems, germination is induced at the start of a relatively short three to four month winter rainy season, and the majority of plants grow well through this rainy season, and only complete their life cycle during the subsequent period when the water is being depleted (Seabloom *et al*. 2003; Levine & HilleRisLambers 2009). The general decrease in water availability after the end of the rainy period as well as between-species variation in their tolerance to dry conditions results in large interspecific phenological variation (Godoy & Levine 2014; Alexander & Levine 2019). Some species complete their life cycle in spring while others persist through fall. This phenological variation and its association with competitive dynamics have been investigated in several past empirical studies (Godoy & Levine 2014; Kraft et al. 2015; Alexander & Levine 2019), providing key real world tests for the models we develop here.

## Water competition model

We develop a general model of plants competing for water when soil water availability determines the duration of each competitor’s growing season. Although we tailor the model to a community of seasonally water-limited Mediterranean annual plant species that differ in their ability to grow under dry conditions, the findings should apply more broadly. The outline of the model is as follows: Plants grow unfettered by neighbors until neighbors reduce the water content of the soil, *W*, to less than the species-specific minimum water content required for an individual to grow, 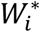 (mass of water per unit land area or volumetric water content with fixed soil depth). Individual species successively stop growing and convert biomass to seeds, and after the last species does so, the season begins again. In this section we build a quantitative model which formalizes these dynamics.

A year begins with all individuals of all species in the seed stage, and all germinate synchronously after the onset of the rainy season. We assume for simplicity that negligible water is carried over in the soil from one year to the next, that negligible water is lost to run-off, ground water and evaporation from the soil (not transpiration), and that rain ceases on the day plants germinate. All of these simplifying assumptions can be relaxed without changing the qualitative results.

After germination, plants grow at a constant rate (when time is measured on a nonlinear scale) until the soil water drops below the species-specific critical value. Specifically, we assume that the biomass and crown area of an individual are related by a power-law allometry with exponents common to all species. When a plant is actively growing, its rate of carbon gain is proportional to its crown area, which implies that plants do not shade one another (Appendix 1.1). This property, together with the power law allometry imply that, at time *t* during the growing season, a species-*i* individual’s biomass is approximately (*G_i_t*)^*B*^, where *G_i_* is a species-specific constant growth rate and *B* is a constant common to all species (Appendix 2). Therefore, if we use the nonlinear time scale *τ* = *t^B^* in place of ordinal time *t*, individuals of species *i* grow at a constant species-specific rate, 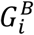, until the time at which soil water content drops below that species’ critical value 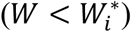. We denote this time *t_i_*, or in the nonlinear time scale *τ_i_* (which equals 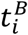). Also at this time all individuals of species *i* convert their biomass 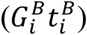 to seeds following a proportionality constant shared across all species, and these seeds survive to initiate next year’s growth process.

Competition emerges in this model only because individuals uptake and transpire the limited soil water pool, and thereby shorten the growing season lengths of competing (or species). The greater the abundance of conspecific or heterospecific competitors, the faster the uptake and depletion of the finite soil water pool, and the earlier soil water drops below each species’ critical value. A plant’s biomass accumulation rate during the period suitable for its growth, 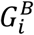, is unaffected by competition from other individuals (Box 1); only the duration of growth is affected by competition.

Three seemingly special assumptions make competition for water in this system analytically tractable – (1) plants have constant growth until a season’s end determined by a soil water threshold, (2) competition between species shortens growing season lengths without changing growth rates during the growing season, and (3) end-of-season fecundity is proportional to 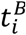, with B common to all species. Importantly, all are emergent properties of plant physiology under a simple hydraulic model (described in Box 1 and Appendix 1, 2) provided that species adhere to a power-law allometry. We both empirically verify this allometry and justify it theoretically in Appendix 2.1, 2.2, and 2.4.

### A single species system

In this section, we quantitatively describe the dynamics of a single species in a water limited system, because the multispecies model follows naturally from this simpler case. To keep the notation simple, in this section we omit the species-specific subscript from all quantities *except the season length τ*_1_, to distinguish it from the within-year time *τ*. Let *N_T_* be the population density of seeds in the soil starting year *T* (note: upper case “*T*” designates year, whereas lower case “*t*” or “*τ*” designates time since germination within a year). Also, because the growing season length is a decreasing function of the population size, we write it as *τ*_1_(*N_T_*). We assume the number of seeds available to start the season is proportional to the prior year’s end-of-season biomass according to the constant *F*. This yields the following expression for per capita population growth in year *T*

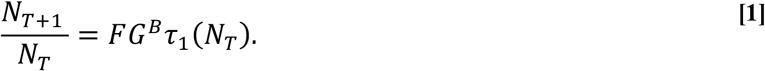

Since both *F* and *G^B^* are constants, the per capita population growth rate of species *i* varies solely as a function of its growing season length, *τ*_1_, which is itself a function of the rate at which water is withdrawn from the common pool.

Given that species compete with one another by shortening their growing season, it is useful to define the length of the season for which the population shows zero growth. Note that the lifetime reproductive success (LRS) of a recently germinated seedling equals the expected number of germinable seeds it will produce during its lifetime, and that this increases with season length, *τ*_1_(*N_T_*). We call the “break-even time” *τ*\* - the value of *τ* at which LRS equals one (or equivalently, *N*_*T*+1_ = *N_T_*). By definition, *τ*_1_(*N_T_*) = *τ*\* at the population dynamic equilibrium. We calculate *τ*\* by setting 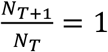 in equation 1:

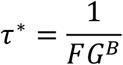

If *τ*_1_(*N_T_*) > *τ*\*, plants outlive the species’ break-even time, making LRS > 1 and causing abundance to increase, whereas if *τ*_1_(*N_T_*) < *τ*\*, the species’ population decreases. This formula truns out to be the same in the multispecies problem described in the next section, where the break-even time is species-specific: 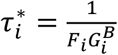.

We now re-write equation 1 in terms of the break-even time *τ*\* as

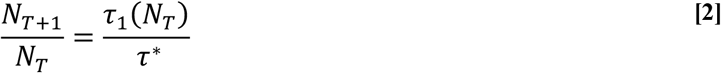

which reveals the direct relationship between growing season length and population growth. The biomass conversion to seed, *F*, and within season biomass growth rate, *G_B_*, remain important determinants of between year population growth, but their effects operate by changing a species’ break even time *τ*\*. A species producing more seeds per unit biomass (higher *F*) can replace itself with less biomass production over the year, and can thus sustain a population while stopping growth earlier in the season (have a lower *τ*\*).

Water availability affects population growth across years via its effect on the length of a species’ growing season *τ*_1_(*N_T_*), which is itself a function of the depletion of soil water. Putting aside for the moment the details of the depletion process, a species can put on biomass as long as the soil water availability exceeds its critical water threshold. This leads naturally to a condition for positive growth when rare. A species can invade an empty system from low density provided the amount of water available at its break-even time, 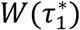, is greater than its critical soil water content, *W*\*, as this implies it is able to outlive its break-even time. Since a species has negligible effect on soil water content at low density, this is true when:

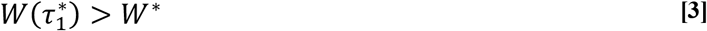

where *W*(*τ*) describes pre-invasion, within-season water availability as a function of *τ*, and 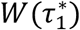 gives the value of this function at the species’ break-even time. We denote the amount of water available at the beginning of the growing season *W*_0_, and in a system with no plants, soil water stays at this level through the entire season *W*(*τ*) = *W*_0_. What follows is a very simple graphical method for determining whether a species can invade the system with no other competitors present. First, we plot water availability through the year, *W*(*τ*) = *W*_0_ on a plot of *W* versus *τ*, and then add a point corresponding to the species’ critical water content, *W*\*, and break-even time, 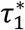, (Fig. 1A). If that point lies above the water availability line, *W*\* > *W*_0_, the water supplied is less than the species’ soil water threshold and the population goes extinct (Fig. 1A). If the point lies below the water availability line, then *W*\* < *W*_0_, more water is supplied than the species’ threshold water level, and the species will successfully invade (Fig. 1A).

**Figure 1:**
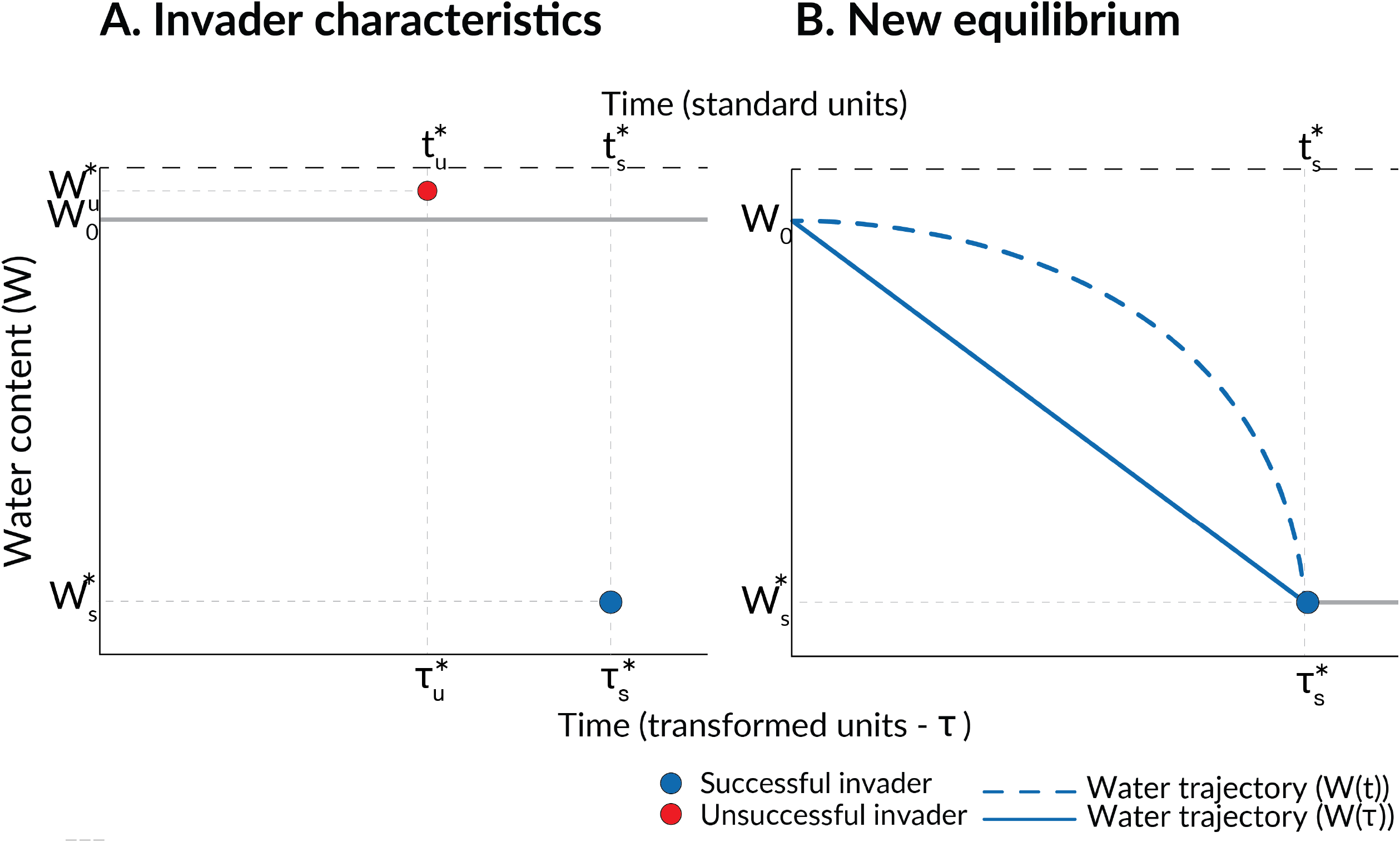
Graphical method for determining invasion success for a single-species system. In Panel A, the gray line denotes the pre-invasion water trajectory, *W*(*τ*), which remains flat at the initial water content, *W*_0_, due to the lack of evapotranspiration, evaporation or precipitation. The red dot denotes the characteristics of an unsuccessful invading species with break-even time 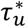(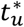 in standard units) and critical soil water content 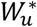. The invader is unsuccessful because its critical soil water content is greater than initial water content, and therefore cannot increase in biomass. The blue dot denotes the characteristics of a successful invading species with break-even time 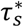(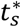 in standard units) and critical soil water content 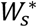. The subscript ( denotes a successful invader, while the subscript *u* denotes an unsuccessful invader. In Panel B, the characteristics of the successful invader are again denoted by the blue dot. In the resulting single-species equilibrium, the solid blue line describes the water drawdown trajectory, *W*(*τ*), from the initial water content to the invading species’ critical soil water content at *τ* = *τ*\* in transformed time units. The dashed blue line shows the water drawdown trajectory in standard time units, *W*(*t*).

To determine a species’ population growth rate 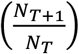 and growing season length (*τ*_1_(*N_T_*)) away from equilibrium, as well as the resulting soil water trajectory, *W*(*τ*), we must model water depletion. The water available at the end of the species’ growing season, or equivalently, that species’ shutoff point, *W*\*, equals the water available at the beginning of the growing season, *W*_0_, minus the water consumed by the species from the beginning of the season to the end, *τ*_1_(*N_T_*). More formally:

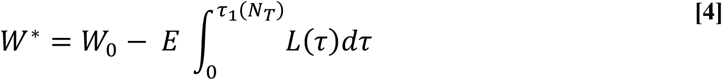

where *E* is the evapotranspiration rate per unit leaf area (Box 1) and *L*(*τ* \\\) is the total leaf area at time *τ* (see Appendix 2.3 for the functional form of *L*(*τ*)). Water consumption is thus a time integrated function of the leaf area of the population from the start of the season to the time its stops growing. Given that *W*\* and *W*_0_ are fixed parameters of the species and system, growing season length is calculated by integrating the right hand side of equation 4 and solving for *τ*_1_(*N_T_*) (see Appendices 3.1 and 5 for the solutions). This value, when placed into equation 2 gives the population growth rate.

Integrating equation 4 and subsequent algebra (Appendix 3.4) produces the within-season water availability, a piecewise linear function of *τ*. Specifically, when *T* > 0:

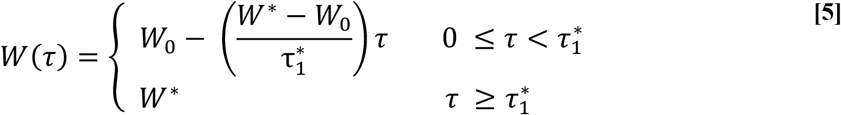

This means that in all years (*T* > 0), within-season water availability, *W*(*τ*), follows a straight line on a plot of *τ* versus *W* from (*τ* = 0, *W* = *W*_0_) to (*τ* = *τ*\*, *W* = *W*\*) as shown in Fig. 1B, which also implies that the species reaches its population dynamic equilibrium in a single year (see Appendix 3.4). The fact that the population snaps to equilibrium in a single year is an artifact of the simplifying assumption that water is only lost from the system by transpiration. Thus, a solitary individual will grow till the water availability drops to its species-specific critical threshold, and at this point its biomass equals what would be achieved by any size population depleting that same water pool. The single year snap to equilibrium does not occur in cases with two or more species.

### A model of two species

The mechanisms governing competition and coexistence of two species can be extended from the rules for a single species to invade. We now label the species using the subscripts 1 and 2 such that species 1 finishes its growing season before species 2 (i.e., 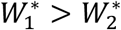). Coexistence can be predicted from just the break-even times and the critical water values of the two competitors. To show this, we take a mutual invasion approach to coexistence, first considering whether an early species can invade a late season competitor. This can be determined by applying the graphical method developed in the single-species case. In fact, identical to the water drawdown in the single species model (Fig. 1B), is the pre-invasion water drawdown, *W*(*τ*), by the late season resident species at equilibrium (solid blue line in Fig. 2A). Here, *W*(*τ*) is given by equation 5 (with subscripts changed to reflect that the species 2 is the resident). The question for invasibility is whether the resident species consumes water too fast for the invading species to reach its break-even time. To answer this, we determine whether 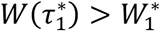 by plotting the point corresponding to the invader’s critical water content and break-even time 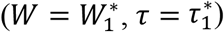. If this point lies above the line corresponding to the water dynamics when the resident is at equilibrium, *W*(*τ*), the invader’s critical water value, 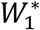, is reached (along the resident’s line) before its break even time 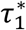, and thus the invasion will be unsuccessful (Fig. 2A). If, instead, the point lies below the water-drawdown line, the system with just the resident species reaches the invader’s critical water value after the invader’s break-even time, and the invasion will be successful (Fig. 2A).

**Figure 2:**
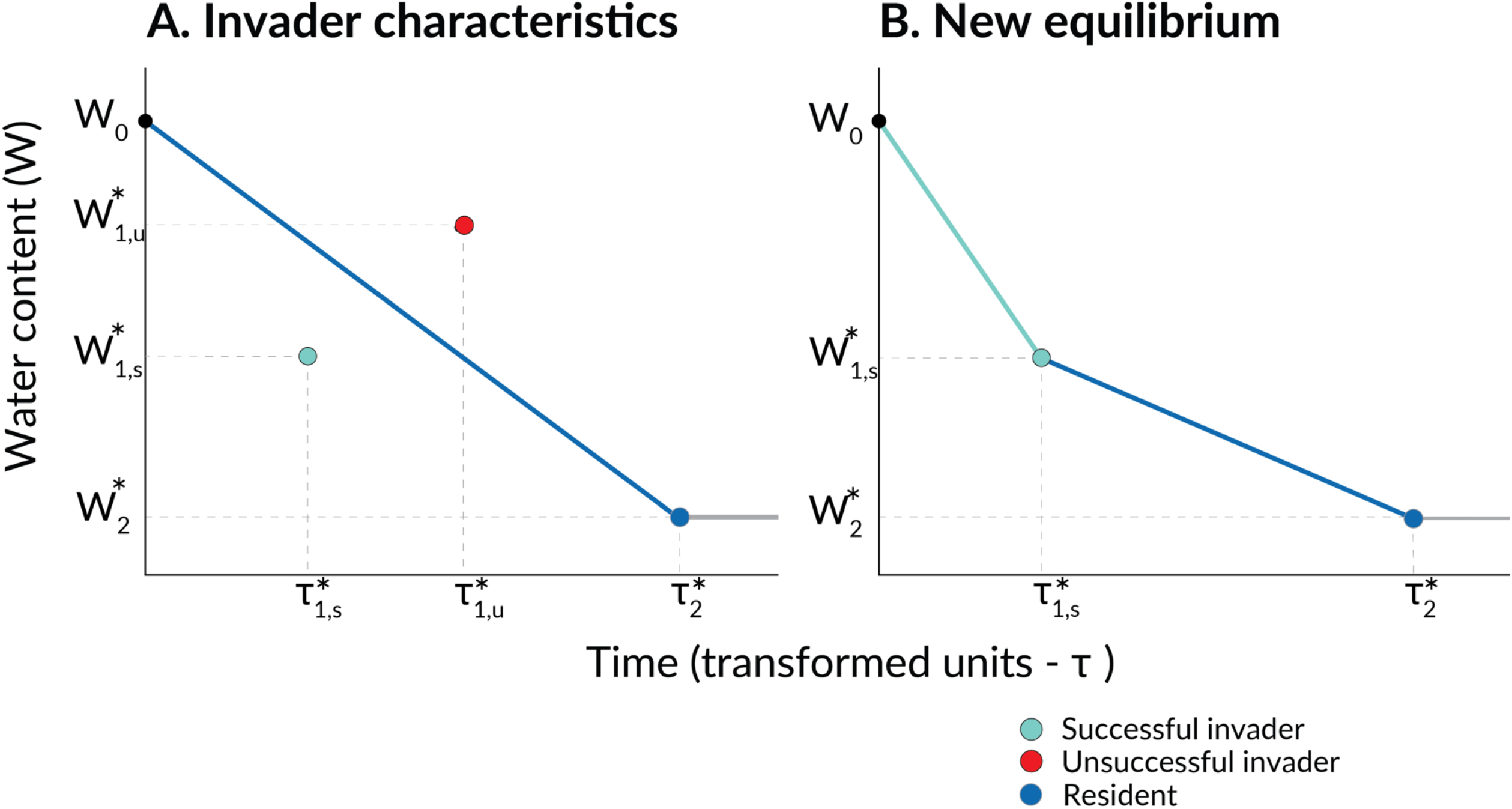
Graphical method for determining invasion success in the two-species system. In Panel A, the late-season resident species (species 2) is the same as the invading species from Panel B of Figure 1, with the equilibrium water drawdown trajectory in transformed time units, *W*(*τ*), shown by the same solid blue line. The characteristics of an unsuccessful early-season invading species (species 1) are denoted by the red dot. Because the invading species’ characteristics lie above the water drawdown trajectory, the resident species depletes water too quickly for the invader to grow to its break-even time. The characteristics of a successful invading species are given by the light blue dot in Panel A. In Panel B, the characteristics of the successful invading species are again shown. At the resulting two-species equilibrium, the water drawdown trajectory, *W*(*τ*), is given by both the light and dark blue lines. In the first phase, in which both species grow, the water decreases steeply to reach species 1’s critical soil water content at its break-even time. Then, after species 1 converts to seed, the water is withdrawn by species 2 alone until its critical soil water content and break-even time are reached.

In contrast to the questionable invasion of an earlier season competitor, a late-season species can always invade an early-season resident species. This is because a late-season species (species 2) has unrivalled access to water after species 1 stops growing, and because we have assumed for simplicity that the only way the soil can lose water is through transpiration. Thus, an infinitesimal invading population of species-2 will grow forever on the finite water available after species-1 has stopped transpiring. It is straightforward to eliminate this unrealistic dynamic by adding evaporative water loss. However, because the results are qualitatively similar, but more analytically cumbersome, we continue to assume a lack of evaporation while recognizing that this does not fully capture the true resource dynamics of the late season.

Given that a late-season species can always invade an early-season resident, the invasion condition for the early-season species, 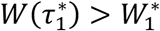, is sufficient for mutual invasion. Substituting equation 5 into this condition, we find:

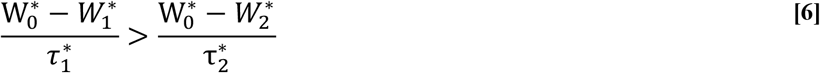

(see Appendix 5.1 for derivation). This implies that in order to coexist, the earlier species must be able consume water faster over the time is takes to produce one new individual per capita than the later species consumes water over the time it takes to produce one new individual per capita. If this condition is met, species 1 is more than compensated for its reduced access to water by a growth rate advantage (shorter 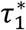) relative to species 2. If species 1 not a sufficiently fast grower during its shorter period of water access, it will be excluded.

If a species is successful in invading the resident, the population density of the invading species will increase, and that of the resident species will decrease until both reach their new, two-species equilibrium population densities (Appendix 3.3). As with the single species model, for all years *T* > 0 the within-season water availability, *W*(*τ*), is a piecewise linear function of *τ*, though it now has three phases. In the first phase, both species grow and consume water until species 1’s critical water content, 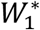, is reached. Then, in the second phase, only the late-season species continues growing and consuming water until its critical water content, 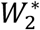, is reached. Finally in the third phase, water availability remains at 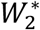 until the season ends. At equilibrium, *W*(*τ*) for a two-species system is given by:

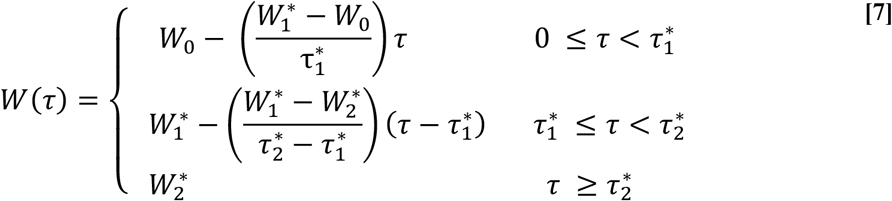

Thereby at equilibrium, water availability follows a straight line on a plot of *τ* vs *W* first from (*τ* = 0, *W* = *W*_0_) to 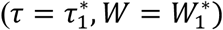, and then from 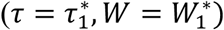 to 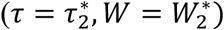 (Fig. 2B).

If the mutual invasion condition is satisfied, it implies the existence of a feasible two species equilibrium that is both unique and globally stable (see Appendix 6). In fact, a time-dependent solution for the population densities of each species can be derived if there are at most two species (Appendix 6). As in the single species case, the equilibrium is maintained because perturbations in either species’ densities are corrected in subsequent years due to the feedbacks between altered water availability, growing season length, and population growth rate. That equilibria are unique and globally stable supports the assertion that mutual invasion is a necessary and sufficient condition for stable coexistence in the model.

### A model of Q species

We now extend the two species result to an arbitrary number of species, *Q*, and label species *i* = 1,…, *Q*, such that species 1 again has the shortest growing season length while species *Q* has the longest (i.e. *W*_1_ > *W*_2_,…, *W_Q_*). We again develop a graphical methodology for determining invasion success for systems with more than two species. Figure 3 illustrates this technique for a two-species resident community (the same one from Figure 2, Panel B) and single invader species with an intermediate critical water threshold. Such an invasion can be successful in a two-species resident community provided the point corresponding to its critical water threshold and break-even time lies below the water drawdown trajectory, *W*(*τ*), for the resident community. This indicates that the resident community consumes water slowly enough for the invader to reach its break-even time before the water level drops to its critical water threshold. (Figure 3).

**Figure 3:**
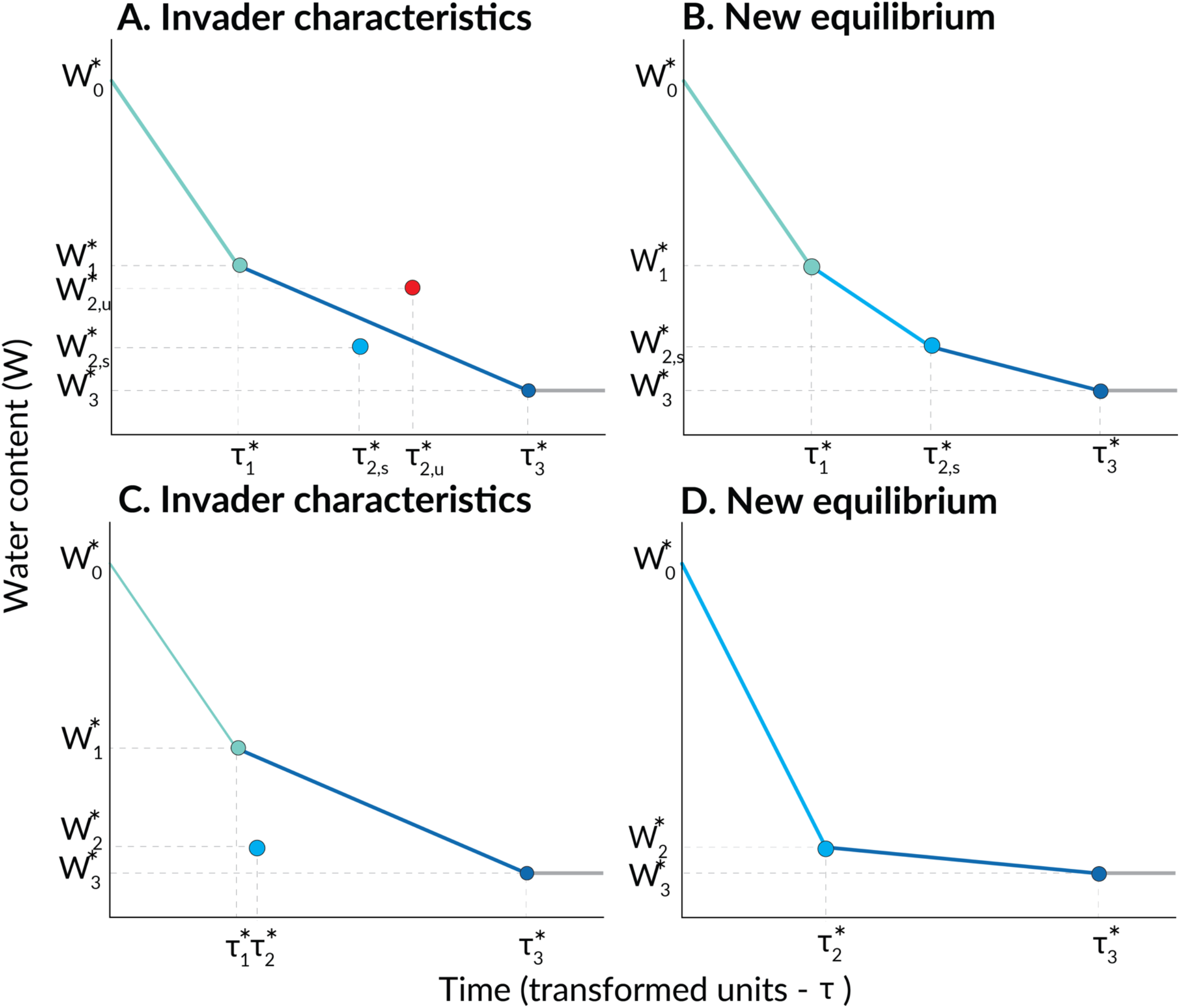
Graphical method for determining invasion success – three-species system. In Panel A, the resident two-species community (species 1 and 3) is the same as that from Panel B of Figure 2, with the equilibrium water drawdown trajectory, *W*(*τ*), shown by the same solid blue lines. The characteristics of an unsuccessful intermediate-phenology invading species (species 2) are again denoted by the red dot. Because the invading species’ characteristics lie above the water drawdown trajectory, the resident community consumes water too quickly for the invader to grow to its break-even time. The characteristics of a successful invading species are given by the blue dot of intermediate hue in Panel A. In Panel B, the characteristics of this successful invading species are again displayed. At the resulting three-species equilibrium, the water drawdown trajectory is given by three solid blue lines. In the first phase, in which all three species grow, the water decreases steeply to reach species 1’s critical soil water content at its break-even time. Then, after species 1 converts to seed, the water is withdrawn by species 2 and 3 until species 2’s critical soil water content and break-even time are reached. Finally, species 3 withdraws water alone until its critical soil water content and break-even time are reached. In Panel C, the characteristics of a successful invader are again shown by the blue dot of intermediate hue. This species will invade successfully, and subsequently cause species 1 to be excluded, as the resulting community will withdraw water too quickly for species 1 to grow to its break-even time. Panel D shows the resulting two-species equilibrium, in which only species 2 and species 3 persist.

At equilibrium, water drawdown in a *Q*-species system is again given by a piecewise linear function of *τ*:

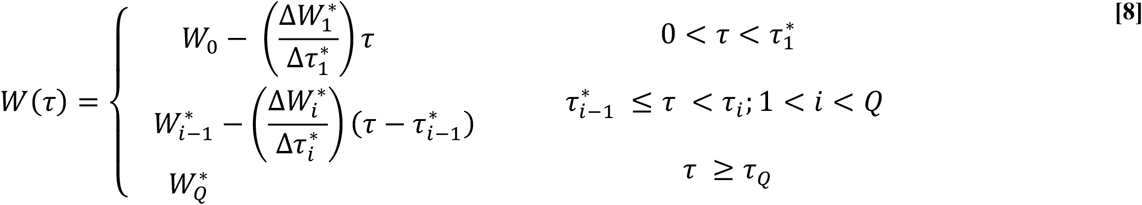

where 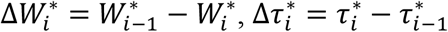, and *τ*_0_ = 0. The growing season is thereby divided into *Q* + 1 periods, each defined by the number of species left growing and consuming water. Species drop out of the system sequentially as their critical water contents are reached, until none remain.

Using the invasion condition, 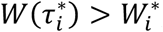, and equation 8, we find that mutual invasion in a *Q*-species system requires that the following expression (see Appendix 4) is satisfied for all adjacent pairs of species, once ordered according to 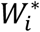:

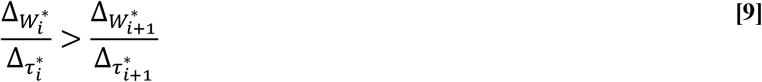

This condition requires that later-phenology species cannot consume water at a rate which prevents earlier-phenology species from reaching their break-even times at low density.

Importantly, a species which successfully invades a Q-species resident community can cause earlier residents to be excluded (a later resident always persists following the reasoning in the two-species section above). Returning to the case of a species with intermediate critical water content invading a two-species resident system: exclusion of the earlier resident will occur if the invading species has a faster water consumption rate at its single-species equilibrium than the earlier resident (i.e. if equation 9 is not satisfied for *i* = 1; Fig 3B). In this case, the invading species has a greater biomass growth advantage (lower 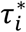) in proportion to its water access than the resident does, and will therefore consume water fast enough that the resident with higher critical water threshold can no longer reach its break-even time at low density (Fig. 3C).

In a system of Q competing species, the subset which eventually coexist can be determined using a simple graphical algorithm. To start, we plot the minimum soil water content, 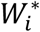, and break-even time, 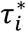, for each of the Q species as we have done in the single, two and three species cases (Figure 4A). Next, we draw a line from the point corresponding to the initial water content (*W* = *W*_0_, *τ* = 0) to the species’ point for which the resulting line has the steepest negative slope. Then, we again draw a line from that species’ point to another species’ point, again minimizing the slope of the resulting line. We continue connecting the dots from left to right until we cannot draw a new line that is negative in slope. Following the condition in equation 9, all of the connected species will coexist, the rest will be excluded. The set of species which persist in equilibrium is thus given by the lower, convex hull of the point cloud that results from plotting each species’ minimum water content and break-even time, as well as the point corresponding to the start of the season (*W* = *W*_0_, *τ* = 0).

**Figure 4:**
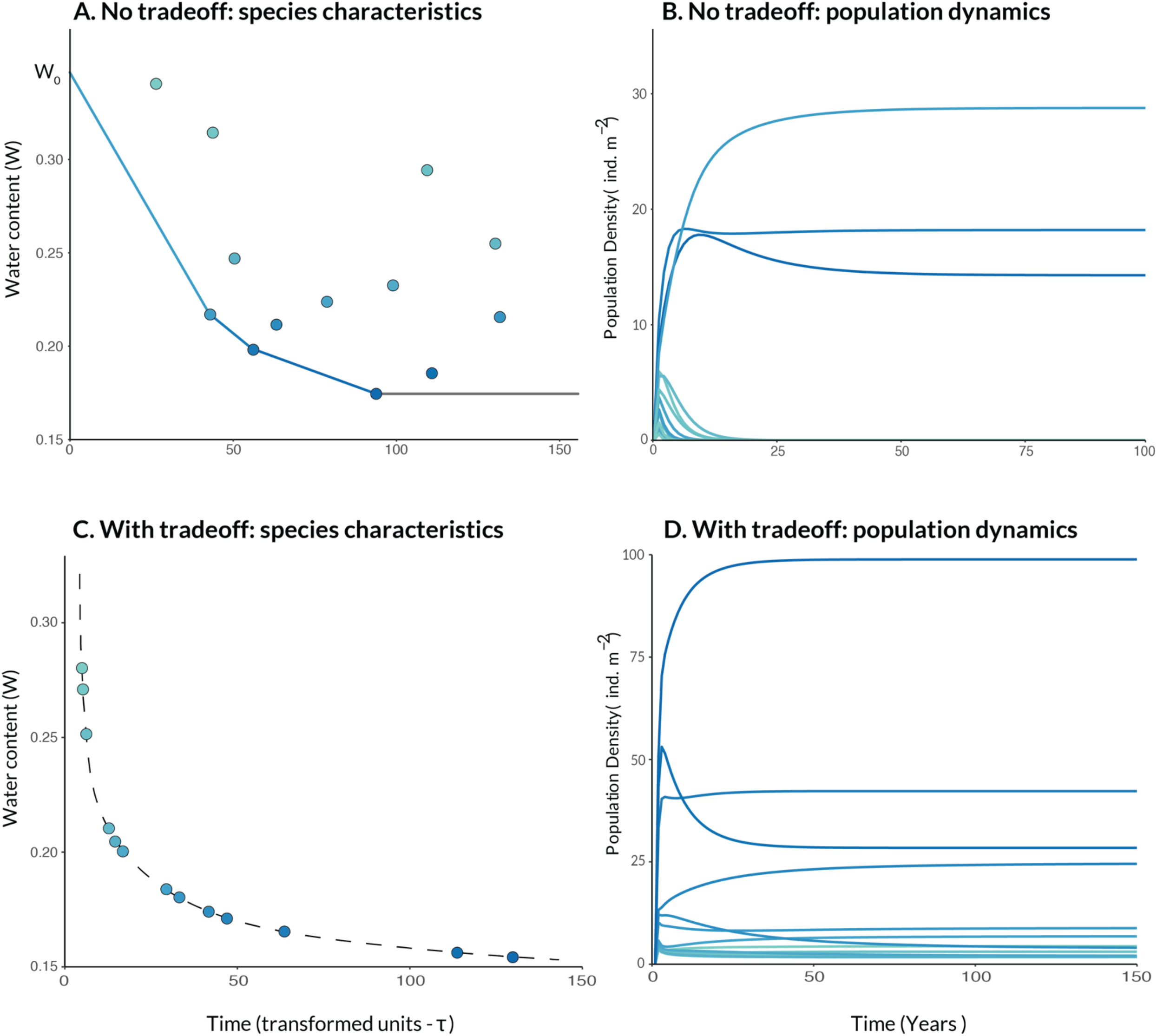
Coexistence outcomes in large species assemblages as a function of an interspecific tradeoff. Panel A shows an assemblage of 13 species with randomly selected critical soil water contents and breakeven times. The graphical method is used to determine the subset of these species which will persist in time. The characteristics of these three species are connected by solid lines which denote the water drawdown trajectory at the resulting equilibrium. Panel B displays the population dynamics in time for the 13 species assemblage, with the three coexisting species forming an equilibrium and the rest collapsing to zero. Panel C shows an assemblage of 13 species with characteristics chosen according to a tradeoff curve which satisfies the coexistence conditions. Panel D displays the population dynamics for these species over time.

An interesting consequence of the coexistence in Fig. 4A is that after community assembly all coexisting species will appear to exhibit a tradeoff, in that the early-season species will have both faster growth and a higher minimum water threshold than the late-season species, even if no physiological tradeoff exists in the species pool prior to assembly. This emphasizes that one cannot infer the existence of a physiological tradeoff from the traits of coexisting species.

If instead of forming a point cloud like in Fig. 4A, species adhere to a strict decreasing and concave-up tradeoff like that in Fig. 4C, then every species in a set that has 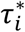 greater than a threshold value (see Appendix 4) will coexist with one another no matter how many are placed in competition. This is again because early-season species are more than compensated for their shortened growing seasons by increased growth rates, allowing them to reach their break-even time before their late-season competitors reduce soil water content below their critical value.

Indeed, there are physiological reasons to expect the very tradeoff allowing coexistence. First note that satisfying the mutual invasion condition in equation 9 requires a tradeoff between a species’ ability to tolerate low water (its 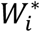), and the speed at which it accumulates biomass (which shortens 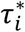). More precisely, for all species to coexist, the functional relationship between species’ critical water threshold and break even time must be negative (higher water thresholds mean faster break even times), and concave up (progressively longer break even times mean lesser advantages in critical water threshold) (Fig. 4). Finally, the tradeoff curve must be restricted to species with minimum water thresholds greater than *W*_0_, because any species with *W*\* > *W*_0_ would go extinct even without the presence of a competitor.

Notably, when allocational constraints are incorporated into the physiological model of plant growth under water limitation, a tradeoff curve that is both negative and concave-up emerges. That model, presented in Appendix 1, and 4.2, predicts that species should be perfectly ordered such that as a species’ growth rate, 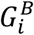, increases (which decreases its 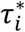), its threshold soil water content necessary for continued growth, 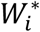, also increases (Box 1, Appendix 1.1). The reason for this relationship is that plant traits that allow transpiration at low levels of soil moisture, such as dense stem tissue that resists embolism (Rosner 2017), require photosynthate that could have been used for productive tissues like leaves. Allocating photosynthate to embolism-resistant structures then slows the rate of biomass growth. The positive correlation between 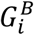 and 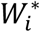, together with the monotonic drying of the soil after the winter rains implies a tradeoff between biomass growth rate and growing season length, 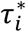, that has negative slope. As we explain more fully in Appendix 4.2, the tradeoff is also likely to have a concave-up shape because the growth cost to reduce one’s critical soil water content increases in magnitude for lower values of *W*\*. These accelerating growth costs mean that as *W*\* decreases, the corresponding changes in break-even time (the inverse of growth) also increase in magnitude (Appendix 4.2).

Thus far, we have derived expressions for the population dynamics, and invasion conditions for any Q-species system of plants in which each year starts with the same initial level of soil moisture (i.e. saturated soil), which then progressively dries because of transpiration until every plant is unable to grow, and introduced a graphical method that allows one to determine, for any set of species, which will persist if all are placed in competition with one another. In the Appendices, we additionally prove that a Q-species equilibrium, if it exists, is unique (Appendix 3.3). We show that for any combination of species, invasion when rare implies the existence of a positive equilibrium (Appendix 4.1). Moreover, we develop a complete understanding of the two-species system, including a time-dependent solution for species abundances, and prove that in such a system any positive equilibrium is globally stable (Appendix 6). We also prove that in a Q-species system any positive equilibrium is locally stable, which is consistent with the fact that a positive Q-species equilibrium implies the successful invasion of each of the Q species into an equilibrium community of the remaining Q-1 species, and that each of these Q-1 species equilibria is also positive (Appendix 7). Last, we showed that the tradeoffs which imply the existence of a stable equilibrium in the population dynamics correspond precisely to a model of plant physiology (Appendix 4.2), suggesting that our coexistence criteria may be generally satisfied for annual plant communities whose physiology is accurately described by our model.

#### Empirical system and qualitative model validation

Our prior empirical work with phenologically distinct annual plants at the University of California, Sedgewick Natural Reserve in northern Santa Barbara County, California, USA, (Godoy & Levine 2014; Alexander & Levine 2019) has identified several qualitative patterns that our model should be able to reproduce. First, a tradeoff should exist such that species that stop growing earlier in the season grow faster than later taxa over their period of overlap. Second, competitive exclusion is always suffered by the species with the earlier phenology; the later species persists due to its access to late season water. Third, earlier phenology of a late season species increases its competitive effect on earlier taxa and reduces their invasion growth rate.

First, we evaluate the tradeoff. Godoy and Levine (2014) quantified the biomass accumulation of three non-native plant species in the system, *Bromus madritensis, Centaurea melitensis*, and *Lactuca serriola*, as well as the date they stopped growing. These taxa were in fact chosen for study because they stop growing at different times of the year. Not only do these species exhibit something close to power law growth, as assumed here, but each species is the most rapid accumulator of biomass during their period of overlap with later competitors (Fig. 5A). This result, of course, is consistent with tradeoffs that underlie coexistence in our model. Assemblages of early, middle, and late *native* species in the same study did not show such tradeoffs, but it is unclear whether these taxa would coexist with one another at the local scale of the study (Kraft *et al*. 2015).

**Figure 5:**
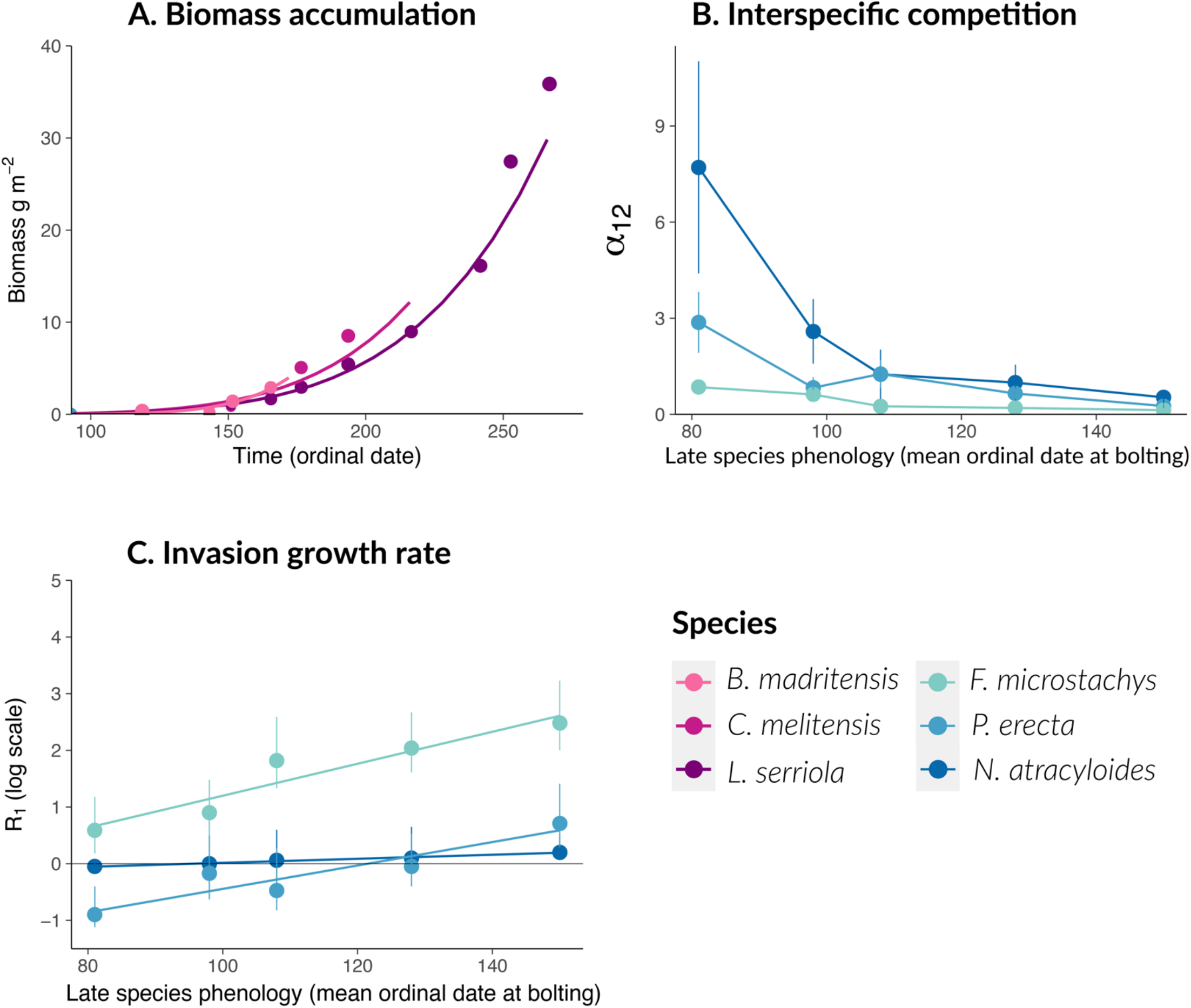
Patterns from empirical studies of phenology and coexistence in California Mediterranean annual plant communities and predicted patterns from theoretical model. Panel A shows the change in biomass over time for three species of differing phenology (*B. madritensis, C. melitensis*, and *L. serriola*). Empirical data is denoted by the points, colored according to species phenology. Power-law fits to the data are shown as solid lines. Panel B shows the per capita competitive effect of a late season species, *L. serriola*, on three resident species of varying phenology (*P. erecta*; *F. microstachys*, and *N. atractyloides*) (*α*_12_) as a function of late-season species phenology, expressed as the mean ordinal date at bolting. Panel C shows the log low density growth rate (*R*_1_) of three resident species of varying phenology (*P. erecta*; *F. microstachys*, and *N. atractyloides*) as a function of the phenology of a late season species, *L. serriola*. In panels B and C, error bars describe the variation around mean values (points). In panel C linear regressions are overlaid to show the trend.

Second we evaluate the pattern of competitive exclusion. Godoy and Levine (2014) parameterized Lotka-Volterra type population dynamic models of competition between each native community (early, middle, and late), and each non-native species (Fig. 5B). These models predicted competitive exclusion for 8 of 9 pairs. Importantly, regardless of the species’ native or non-native status, it was always the earlier species that was excluded. This qualitative pattern is consistent with results from our model, where exclusion always eliminates the earlier species because the later competitor has unique access to water unconsumed by its earlier counterpart.

Third, we evaluate our model’s ability to predict the competitive effect of a later taxa as a function of its phenology. Alexander and Levine (2018) explored how differences in phenology between populations of a late season invader, *Lactuca serriola*, affected its suppression of earlier phenology native plants. Results showed that earlier *Lactuca* populations exerted greater per capita competitive effects on the per germinant fecundity of native plants (Fig. 5B), which reduced their projected invasion growth rates (Fig 5C). Both of these results also emerge from our model. To see how, first note that the per capita competitive effect of a later competitor on the growth of an earlier one is calculated as 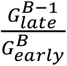 (Appendix 3.6). From this expression, it is clear that a higher biomass growth rate of the later competitor will increase its competitive effect. A higher biomass growth rate of earlier *Lactuca* would be expected if the species-level tradeoff between phenology and biomass growth rate (Fig. 5A) holds within species. This same mechanism in our model also causes an earlier species invasion growth rate to decrease as the later species phenology moves up in time, matching the empirical data (Figure 5C).

## Discussion

In this paper we introduced a mechanistic model of phenologically structured water-competition between annual plants. Our model is built on a foundation of realistic ecohydrology, plant physiology and allometry, can readily replicate and explain empirically observed patterns, and provides several fundamental insights into the interactions allowing for the coexistence of annual plants competing for water in Mediterranean climates. In the model, phenological differentiation arises naturally through interspecific variation in the critical soil water threshold and an evapotranspiration-driven, monotonic decrease in water content through the growing season. When accompanied by a tradeoff in growth rate, the resulting phenological nesting of competitors can maintain infinite species diversity under simple but realistic assumptions.

### The mechanism of coexistence

Pairwise coexistence occurs in the model because an earlier competitor with a higher water threshold, if it grows fast enough, achieves the biomass necessary for positive population growth before all its later competitors deplete the water to its critical value. Meanwhile a later species always persists with earlier, more water demanding taxa due to its ability to grow during the period after these earlier taxa senesce. Moreover, this extends to any number of earlier and later competitors. In a Q-species system the growing season is subdivided into periods with stepwise decreases in diversity as a result of interspecific differences in critical soil water content and a monotonically decreasing soil water pool. When this pattern is paired with a relationship between species’ break even times and critical water thresholds that is both negatively sloped and concave-up (Figure 4, Panels C and D), each species is the dominant competitor for water in exactly one time period – the one in which they are the earliest-phenology species still growing. Under such conditions, each species is able to grow fast enough to achieve positive population growth before its competitors reduce water to its critical value.

Whether or not the appropriate ecophysiological tradeoff between break-even time and critical water threshold exists, any set of coexisting species will appear to exhibit such a tradeoff simply because it is a requirement for coexistence (Fig. 4A). Species that fall above the tradeoff curve will simply have been excluded. This illustrates how the presence of an apparent tradeoff in traits among co-occurring species need not imply the existence of a true allocational tradeoff related to constraints on how plants are built. To determine whether such a tradeoff exists, one must examine the underlying ecophysiological relationships between traits in a broader pool of species, including those that coexist and those that exclude one another.

A key element of annual plant life history is in part responsible for the high degree of coexistence possible in the model: the invulnerability of the seed stage to competition. Species escape the negative effects of the unfavorable environment created by their competitors by converting their biomass to dormant seeds when they reach their critical water threshold. Though species cannot grow during the portion of the season in which soil water content is below their critical threshold, they do not suffer the negative consequences of competition-induced water limitation during that period. And, of course, the later competitor suffers nothing from the dormant seeds.

### Competition for time

When competitors in the model deplete soil water, they alter the length of each species’ growing season, which in turn determines their invasion success, equilibrium abundance and population growth rate. Thus by differentially depleting a shared water resource, species are effectively competing for time. Competition for time dynamics have also emerged in other physiologicaly grounded resource-competition models. For example, Detto et al. (2021) develop a model of light-competition in both forest and annual plant systems in which species grow unfettered by neighbors until they are overtopped by taller competitors. As in our model, the ability of a species to invade a system of competitors is a function of the amount of time it has to grow before being overtopped. Thus, though species in Detto et al. (2021)’s model ostensibly compete for light, they in fact compete by shortening the growing season lengths of shorter species. Detto et al. (2021) also show infinite coexistence in their model, and this requires a tradeoff between growth duration in form of overtopping ability and growth rate that is similar to the tradeoff required for coexistence in our model. The parallels between these two models suggest there may be something fundamental about temporal competition and compensating tradeoffs that enable many species to coexist.

### How simple dynamics emerge from complex physiology

We find it remarkable how the physiological, population dynamic and hydrologic complexity in this model condenses into the simple graphical algorithm illustrated in Figure 4A. As the Appendices make clear, there are many physiological and individual-level parameters that collectively determine the two critical species-specific quantities 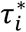 and 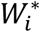. All these parameters can be species-specific with the sole exception of the allometric exponent *B*, which must be common to all species for there to be a single time scale, *τ* that makes the within-season water dynamics linear. And yet, all of this species-specific complexity reduces to just two critical quantities. One implication of this reduction is that incorporating between species variation in other functional traits into the model should not reduce capacity for coexistence, as it will also condense into the two critical species-specific quantities. A second implication is that one should not expect functional trait variation below the level of these two critical quantities to predict coexistence, and this counters to a large literature that attempts to draw such connections (e.g. Weiher & Keddy 2001; McGill *et al*. 2006; Kraft *et al*. 2015).

The simple, tractable dynamics in the model emerge primarily due to the special, but physiologically derived, scaling between leaf area and biomass. In the model, water availability through time is a time-integrated function of the evapotranspiration rate – which is itself proportional to a plant’s leaf area. Reproduction, however, is simply proportional to a species’ end-of-season biomass. Because of the power-law allometric scaling between leaf area and biomass (Appendix 2), leaf area, when integrated, has the same exponent, *B*, as biomass (Appendix 2.3). Thus both evapotranspiration and biomass accumulation, and therefore water availability and reproduction, are linear in the same transformed time scale. This allows the simple graphical method for determining coexistence outcomes, the derivation of expressions for the equilibrium abundances and invasion growth rates for systems with an arbitrary number of species, and the clear stability properties of the model.

### Connection to empirical systems and other coexistence studies

There is substantial evidence that tradeoffs between phenology (degree of tolerance to dry soil) and biomass growth rate of the sort required for coexistence in our model are common in natural systems. For example, past empirical studies demonstrate such a relationship in Mediterranean annuals (Angert et al. 2009; Figure 5, Panel A), and other taxa (Ackerly 2004; de Guzman *et al*. 2017). As discussed previously, these tradeoffs may arise either from underlying physiological constraints or simply as a result of competitive sorting. However, there is reason to expect physiological mechanisms involving variation in stem tissue density (and thus in xylem resistance to embolism) and/or root depth to lead to these tradeoffs. For example, correlations between stem-tissue/wood density and resistance to embolism have been widely observed across taxa (Hacke *et al*. 2001; Rosner 2017). Similarly, deep roots are more expensive to grow and maintain but allow plants to access water only available late in the season (Schenk & Jackson 2002). Correlations between rooting depth and phenology have been described extensively in Mediterranean annual systems (Gulmon *et al*. 1983; Godoy & Levine 2014; Kraft *et al*. 2015), and other systems and taxa (for example, Wright & van Schaikt 1994). Incorporating allocational tradeoffs between rooting depth/stem-tissue density and growth rate into the physiological model developed in Appendix 1 results in a tradeoff curve that is negative and concave-up, as required for infinite coexistence.

Several other studies have examined the coexistence dynamics of plants competing for water. For example, several studies develop empirically validated models of water competition between deep-rooted tree and shallow-rooted grass species to explore Walter’s two-layer hypothesis for tree-grass coexistence in dry savannahs (Walter 1971) (Walker *et al*. 1981; Eagleson & Segarra 1985; Ward *et al*. 2012). The results of these studies are similar to our own, in that coexistence is possible because species with reduced access to water-the grasses (or early-season annuals in our study), are compensated with greater competitive ability. However, while these studies recognize the importance of a tradeoff in maintaining coexistence, they do not consider systems with more than two strategies.

### Limitations

Our model’s tractability results from several key simplifications of nature. The first of these is the approximation of growth as a step function of water availability (Appendix 1.3). This approximation results in a negligible overestimation of total within-season growth relative to the slightly more continuous function predicted by ecophysiology. This difference is unlikely to significantly change our results due to both its small size and consistency across species. A second major assumption is spatial homogeneity in water availability. In nature, water resources are spatially structured, particularly with depth (Gulmon *et al*. 1983; Seabloom *et al*. 2003), and plants place roots at different depths. In fact, this variation could provide a mechanistic explanation for the phenological differences in our model, as deeper rooted species are able to access deeper pools of water and therefore persist later into the season.

Finally, in our model we assume that initial water availability, *W*_0_, is consistent across years. In Mediterranean systems, however, there is considerable interannual variability in rainfall which would translate to variation in *W*_0_ across years (Heady *et al*. 1977; Schonher & Nicholson 1988; Haston & Michaelsen 1997). While we do not explicitly consider such variability in our analysis, it would not qualitatively change the results of this paper. Variation in *W*_0_ could cause early-season species to be competitively excluded. However, though the identity of the earliest species in an assemblage might change, an arbitrary number of species could still coexist.

### Future directions

Future work could explore competitive dynamics in systems with more complexity by incorporating explicit variation in rooting depth and by examining plants with more complex life histories than annuals. Many studies have explored the impact of rooting depth on competitive ability and coexistence dynamics in plants (Fargione & Tilman 2005; Nippert & Knapp 2007; Ward *et al*. 2012; Holdo 2013; Yu & D’Odorico 2015). However, few incorporate realistic models of abiotic soil water movement, a process with complex (Broadbridge et al. 2017), but potentially important implications for competitive outcomes (Manoli *et al*. 2017). Future efforts could also explore systems with more complexity than annuals. Though models of perennial species may be intractable, simulations and numerical analyses could be used to explore whether tradeoffs between water access and growth rate also allow coexistence in those systems.

Finally, our model generates several, empirically testable predictions about the dynamics of water competition in Mediterranean annual plant assemblages. These include 1) plants grow unfettered by competition until reaching a species-specific, minimum water requirement, 2) increased competition reduces a plant’s fecundity by shortening its growing season, and 3) an interspecific, ecophysiological tradeoff between growth rate and minimum water requirement facilitates coexistence. These predictions could be tested by combining pairwise competitive experiments (Godoy & Levine 2014 and Alexander & Levine 2019), which generate competitive outcomes, with measurements of seasonal biomass accumulation rates and coincident water depletion. Combining such empirical approaches with the type of theory developed here offers the chance to rigorously evaluate some ecophysiological mechanisms underlying plant coexistence in nature.

## Acknowledgements

We thank members of the Levine and Pacala labs for helpful feedback on earlier versions of the manuscript, and Eva Phillips for her advice on figure design. We also thank the University of California, Sedgwick Natural Reserve for their continued support. Finally, this material is based upon work supported by the National Science Foundation Graduate Research Fellowship under Awards #DGE-2039656 and #DGE-2039656. Any opinions, findings, and conclusions or recommendations expressed in this material are those of the author(s) and do not necessarily reflect the views of the National Science Foundation.

## Appendix to: Coexistence in water-limited, phenologically structured annual plant communities

### 1 The physiological model

#### 1.1 Model description

The underlying physiological model describes the basic processes of plant growth and water consumption in Mediterranean annual systems. There are three equations which together comprise the physiological model: one which deals entirely with carbon production via photosythesis, and two which relate the photosynthetic process to plant and soil hydrology.

This first equation is a partially simplified version of Farquhar’s photosynthesis model [Farquhar and Sharkey, 1982], and relates a plant’s carbon assimilation rate (minus root respiration *r_l_*), *a*, to the atmospheric and internal concentrations of carbon (*C_a_* and *C_i_* respectively), and light availability (*V_max_*).

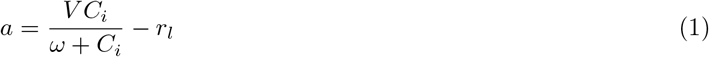

where *ω* is an empirically determined constant. The second pertinent relationship is an adaptation of Fick’s law, and describes the dependence of a plant’s carbon assimilation rate on the plant’s stomatal conductance (*g*) and the gradient of external to internal carbon concentration [Farquhar and Sharkey, 1982].

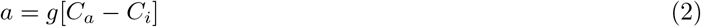

where

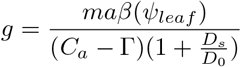

Here, *m* and *D*_0_ are empirical constants, *D_s_* is vapor pressure deficit, Γ is the *CO*_2_ compensation point [LEUNING, 1995], *ψ_leaf_* is the leaf water potential, and *β*(*ψ_l_*) is a shutoff operator bounded by 0 and 1 which goes to 0 as *ψ_leaf_* approaches a plant’s lower physiological limit, *ψ*\* [Wolf et al., 2016]. When the plant is not water limited, *β*(*ψ_l_*) = 1. For convenience, we label the constant portion of *g, ϕ*. Therefore, the above expression simplifies to:

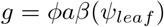

Finally, we use two expressions for evapotranspiration: one which is a function of leaf area and another which is a function of root area. These expressions reflect the fact that evapotranspiration per unit leaf area is dependent on both the combination of a plant’s stomatal conductance, *g*, and vapor pressure deficit, *D_s_*, and the combination of xylem conductivity, *k*, and the water potential gradient from the soil to the leaf. Note that this equation is also a form of Fick’s law.

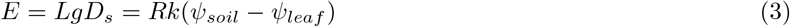

#### 1.2 The carbon assimilation rate

To determine a plant’s growth rate as a function of soil water content, we must first determine the relationship between soil water potential and carbon uptake, *a*. To do so, we first note that when a plant’s leaf water potential, *ψ_leaf_* is above its minimum water potential requirement, *ψ*\*, the plant is not water limited, and therefore operates at the maximum possible carbon assimilation rate, *a_max_*. Once a plant’s leaf water potential reaches its minimum water potential, its stomates begin to close.

We define 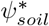 as the soil water potential at which *ψ_leaf_* first reaches *ψ*\*. Therefore, for 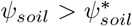:

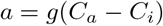

where, recalling that at *a_max_*, *β*(*ψ_leaf_*) = 1:

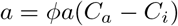

Solving for *C_i_* we find:

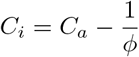

and, plugging this expression into equation 1:

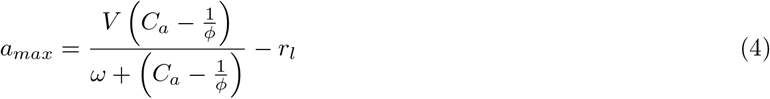

This gives the carbon assimilation rate when the plant is not water limited. Now, to determine 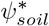, we substitute *ψ*\* and *a_max_* into equation 3, yielding

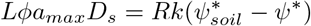

then, solving for *ψ_soil_* we find:

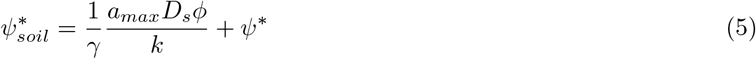

where *γ* is the root-shoot ratio of the plant.

To calculate a plant’s carbon assimilation rate during the period 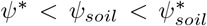, we solve the system of equations 1–3 for a, noting that during this period 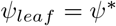.

First, we rearrange equation 1:

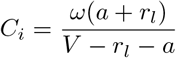

Then, we rearrange equation 3:

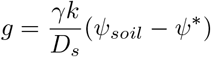

and plug both into equation 2, finding:

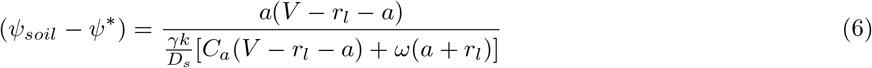

To solve for *a*, we subtract the l.h.s from both sides and find the roots of the quadratic.

#### 1.3 A step-function approximation for the carbon assimilation rate, *a*

The behavior of the physiological model described in section 1.2 prevents tractable analytical treatment for any community of competitors, even in the relatively simple case of Mediterranean annuals. This is because of the nonlinear relationship between *ψ_soil_* and *a* in the period during which *ψ_leaf_* is fixed at *ψ*\*. This nonlinearity prevents us from solving for a species’ single-season reproduction as a simple function of competitor densities. The nonlinear relationships are further complicated by the relationship between volumetric water content, *W*, and soil water potential:

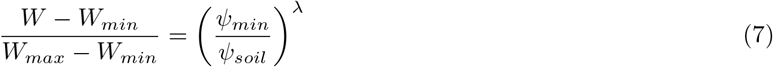

where *W* is the volumetric water content of the soil, *W_max_* is the saturated soil water content, *W_min_* is the residual soil water content, *ψ_min_* is air entry pressure and λ is a dimensionless shape parameter. This functional form comes from the Brooks Corey model [Brooks, 1965], though the van Gnuchten model [Van Genuchten, 1980] is an alternative which performs similarly. The nonlinear relationship between *W* and *ψ_soil_* exagerates the nonlinear portion of the relationship between *W* and *a*, making it more severe, or abrupt. Because of this,equivalent changes in *W* result in much greater reductions in *ψ_soil_* at low values of *W* than high ones. Therefore, at the end of the season when *ψ_soil_* is near *ψ*\*, *ψ_soil_* decreases more rapidly for a given reduction in *W* than at the beginning of the season. As a result, *a* decreases very sharply at the end of the growing season. Further exacerbating this affect is the fact that plant evapotranspiration is proportional to leaf area, which is largest at the end of the season. As a result, *W* declines fastest at the end of the season, when consumption is greatest, accelerating the crash in carbon accumulation rate at the end of the season.

Additionally, there is a positive feedback in the reduction of *a* once *ψ_leaf_* is fixed at *ψ*\*. Carbon accumulation rate, *a*, is a function of stomatal conductance, which is itself a function of *a*. Therefore, any given decrease in *a* results in compounded, larger decreases in the same quantity. In concert with the nonlinear relationship between soil water potential and volumetric soil water content this results in an extremely step-like functional form for the relationship between *W* and *a*. Once *ψ_leaf_* becomes fixed at *ψ*\*, *a* goes to zero very abruptly, rendering the nonlinear period of growth inconsequential.

**Figure 1:**
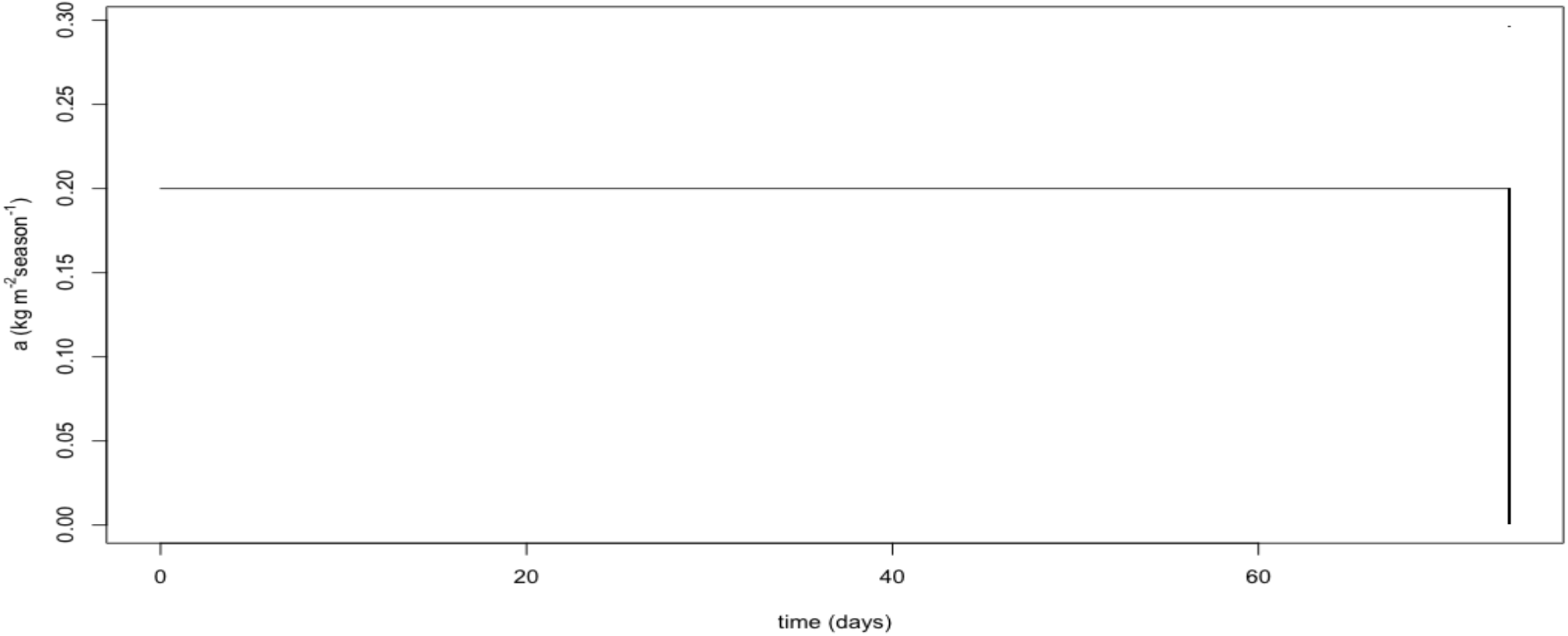
*a* as a function of *time*, with *k* = 0.5*kgm*^−1^ *MPa*^−1^ *season*^−1^

This is best demonstrated by actually calculating and plotting *a* as a function of *W*. Figure 1 shows *a* as a function of *W* with a xylem conductivity of *k* = 2*kgm*^−1^ *MPa*^−1^ *season*^−1^. Choices for the other parameter values were based off empirical estimates from Leuning et al. 1995 and can be found in section 10. With *k* = 2, the relationship is nearly square, exhibiting an imperceptible degree of nonlinearity. This suggests that the relationship between *a* and *W* can be safely approximated as a step function, which in turn allows us to develop a tractable analytical system for competing plants by noting that if *a* is governed by a step-function, plants operate at *a_max_* until the soil reaches *ψ*\*, at which point the plant stops growing for the season and converts its biomass to seed.

To illustrate the robustness of this approximation, we can tweak *k* in an effort to make the function as “nonlinear as possible”. Decreasing *k* to 0.05*kgm*^−1^ *MPa*^−1^ *season*^−1^, an already unrealistically small number, we begin to see a slight degree of curvature in the relationship (Fig. 2)

At the extreme, we can dial *k* all the way down to 0.01*kgm*^−1^ *MPa*^−1^ *season*^−1^, and force some amount of curve into this relationship (Fig. 3). However, even at this unrealistic value, the overestimation of *a* that would occur is slight, especially when one notes that the rate of decrease in *W* is largest in this region given it occurs when plants are near the season-maximal size, and are therefore extracting water much faster than they are at the beginning of the season. Since total growth is a function primarily of time, the error is likely to be small, reinforcing the validity of the assumption.

Figure 4 plots the total estimated carbon accumulation for a plant of given *ψ*\* as well as the total bias resulting from the approximation of the carbon accumulation curve as a step-function for three different values of *k*. The overestimation error is imperceptible in the first two plots and is just a small visible fraction of the total accumulation in the third.

### 2 Allometry

#### 2.1 Determining the appropriate allometric model

To determine the appropriate allometric relationship for Mediterranean annual plants we analyzed a time-series dataset of biomass for 12 species grown at the Sedgewick Natural Reserve in northern Santa Barbara County in California, USA. Data come from [Godoy and Levine, 2014]. We compared two models of the relationship between time and biomass: an exponential model and a power law model. The exponential model is given by:

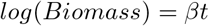

where *t* is the within-season ordinal date, and *β* is an estimated constant. The power law model is given by

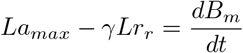

**Figure 2:**
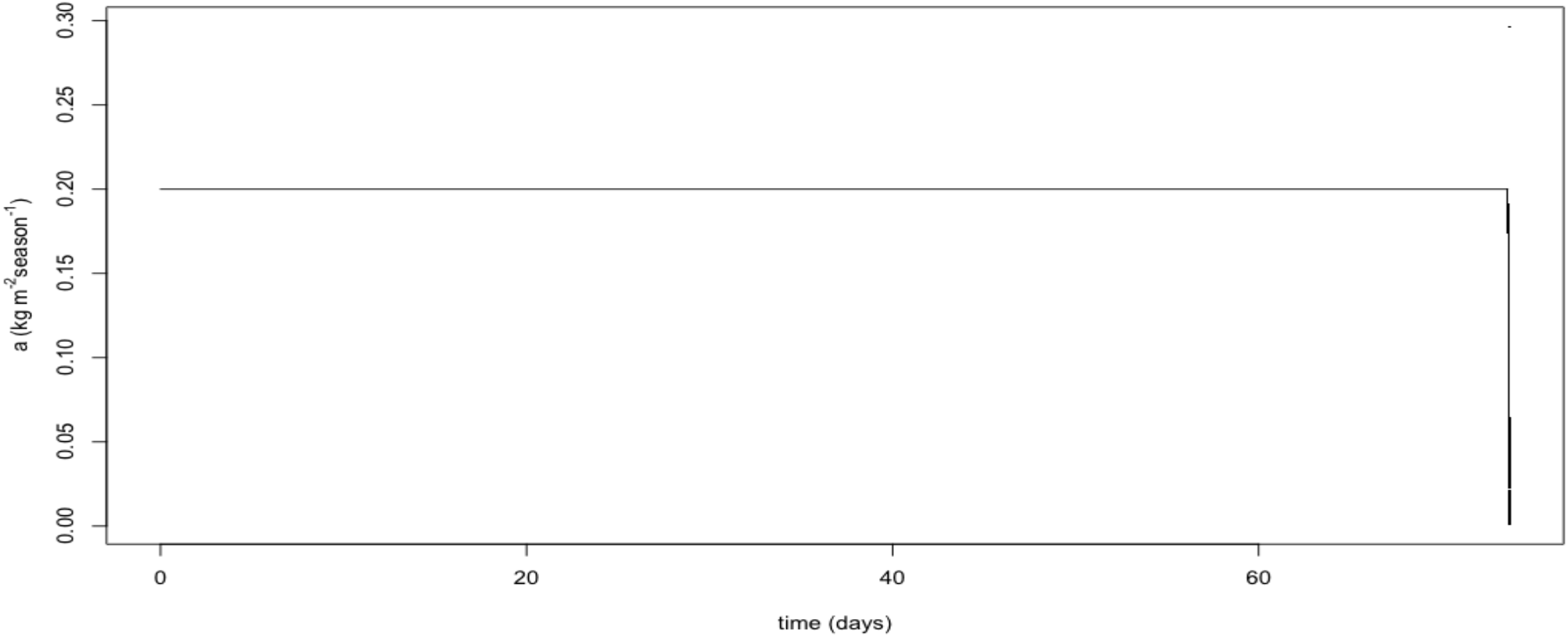
*a* as a function of *time*, with *k* = 0.05*kgm*^−1^ *MPa*^−1^ *season*^−1^

To compare models, we fit both using the lm() function in the R statistical programming environment (R Core Team 2021). Each model includes an interaction effect between time and Species to account for among-species variation in allometric relationships. The power law model exhibited a lower AIC than the exponential model (241.6 vs 289.4), indicating a better fit and that a power-law allometryis more appropriate for the annual plants in our system. Figure 5 shows the observed and estimated relationships from both models.

#### 2.2 Model description

We are primarily interested in two annual plant dimensions: leaf area and biomass. We are interested in leaf area because it is proportional to evapotranspiration, photosynthetic rates, and root area. We are interested in biomass because we assume that a plant’s reproductive output is proportional to biomass by the constant *F*.

Leaf area and biomass are related to each other by the following power law:

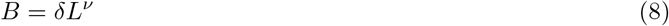

where *δ* and *ν* are empirically determined constants.

#### 2.3 Derivation of the growth rate expression

To calculate a plant’s growth rate, we begin by noting that a plant’s net carbon accumulation rate must equal the change in biomass, *B_m_*, over time

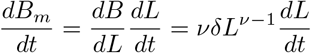

where *r_r_* is the rate of root respiration per unit root area. Recall that we have already accounted for leaf respiration in equation 1. Next we want to solve for the growth rate in leaf area: 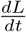. To do so, we first note that

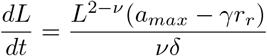

Solving for 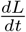 we find:

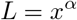

**Figure 3:**
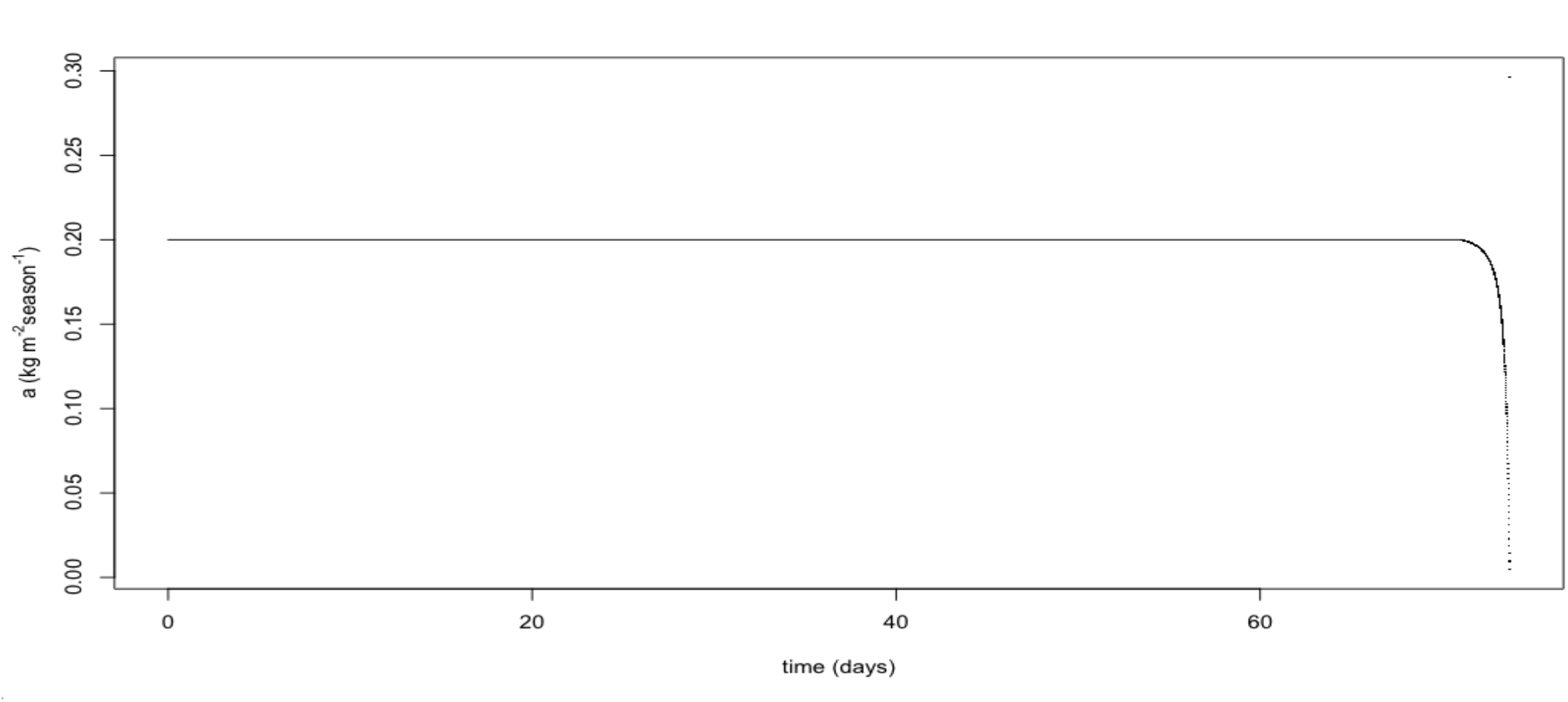
*a* as a function of *time*, with *k* = 0.01*kgm*^−1^ *MPa*^−1^ *season*^−1^

Because we have determined empirically that the allometric relationships are governed by a power law, we can define an arbitraty plant dimension, *x*, in which growth is linear (i.e. 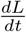 is not a function of *L*). Specifically, *x* is related to leaf area, *L*, in the following manner:

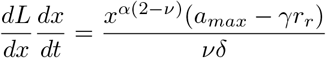

where *α* is an allometric constant. We then plug this expression for L into our expression for 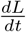:

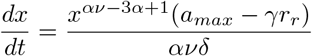

Noting that 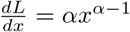, we now solve for 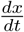, finding:

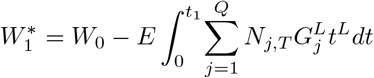

Now, because 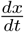 is constant, we can set *αν* − 3*α* + 1 = 0, finding *α* = *L*. This gives us the growth rate in *x*:

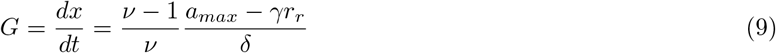

which implies that the leaf area growth rate is:

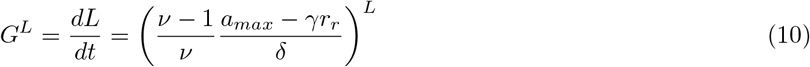

Using this expression we now calculate the total leaf area for species *i* at within-season time *t* and in year *T*:

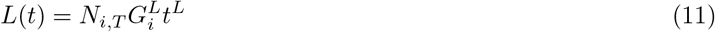

**Figure 4:**
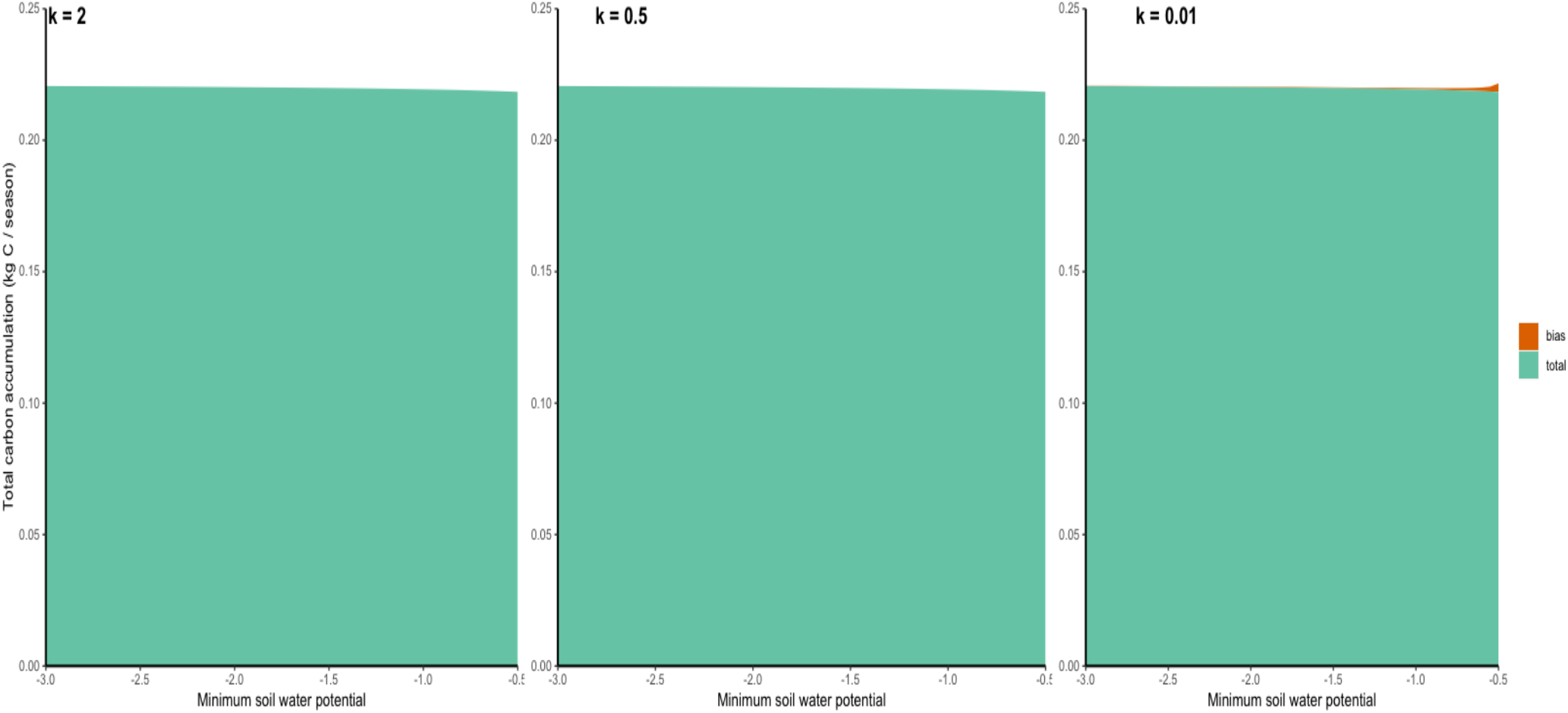
Total carbon accumulation per season as a function of *ψ*\* and total overestimation by the step-function approximation for *a* (in red)

### 3 The full system

The following sections contain full and detailed derivations of the expressions presented and/or referenced in the manuscript. The model notation here is more explicit than that in the manuscript in which constants were aggregated and relabelled for readability. Here, we note where such discrepancies occur and state the equivalent variable in the manuscript.

#### 3.1 Growing season length

Consider a community of *Q* annual plant species which begin growing at the end of the rainy season (*t* = 0) and draw water from a shared pool of soil water as they grow. We label these species from 1 to *Q* such that species 1 has the highest value of *ψ*\*, thus finishes growing first, and species *Q* has the lowest *ψ*_0_, and thus finishes growing last. At the beginning of the season, soil volumetric water content is at field capacity, *W*_0_, which we also refer to as the initial soil water content in the manuscript. Synthesizing equations 3 and 11 we determine

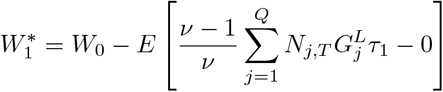

where: *t*_1_ is the time, in standard units, at which species 1 stops growing – *τ*_1_ is itself a function of the population densities of itself and each other competing species, as these competitors reduce its growing season length by consuming water; 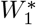 is species 1’s critical soil water content, calculated from *ψ*\* using equation 7; 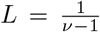; and *E* is equal to *a_max_D_s_ϕ* and gives the rate of evapotranspiration per unit leaf area. Solving the integral we find:

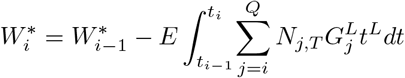

Where 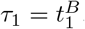. In general we use *τ* to refer to the transformed time scale, *τ* = *t^B^* in which the water dynamics of the system are linear in time. Notice that in the above expression, when *τ* is used in place of *t^B^*, *W* is a linear function of *τ*. Solving for *τ*_1_:

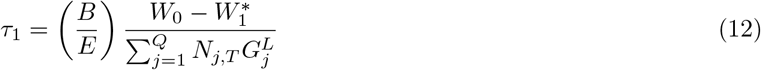

where 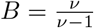, and is equivalent to the exponent in *t^B^* and *GB*. Now, for 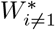 we write:

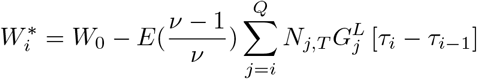

which after taking the integral becomes

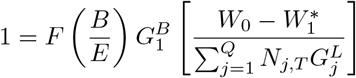

and finally, solving for *τ_i_* gives

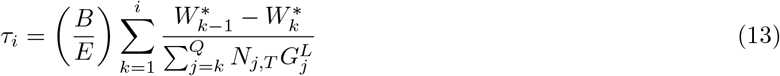

**Figure 5:**
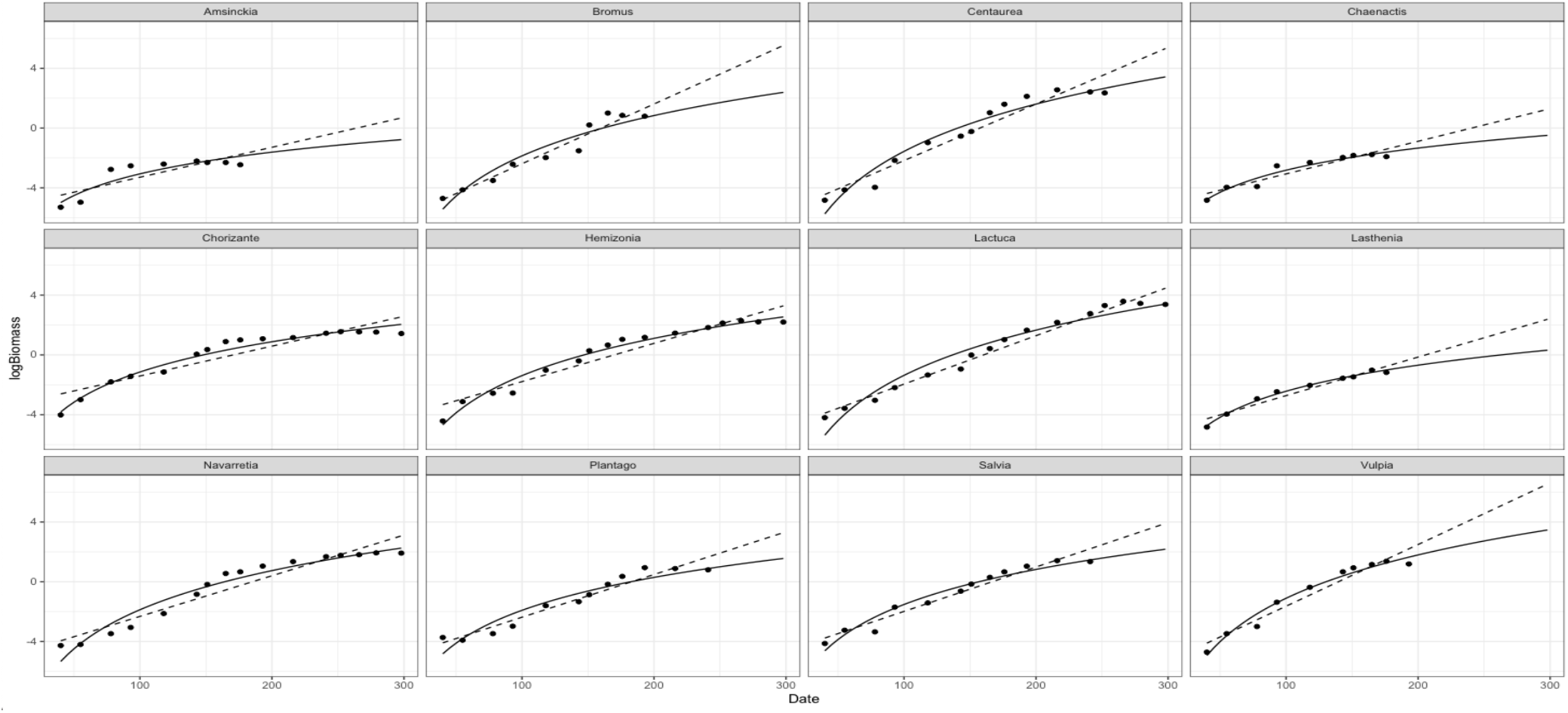
Observed and estimated allometric relationships between time and annual plant biomass. Solid lines represent estimates from the power law model, and dashed lines are estimates from the exponential model

#### 3.2 Population dynamics

Plant reproduction is proportional to biomass by the fecundity constant *F* which is equal to the number of germinating offspring produced per unit biomass. Thereby, using equations 9 and 8 we can write the expression for *N*_*i,T*+1_ as:

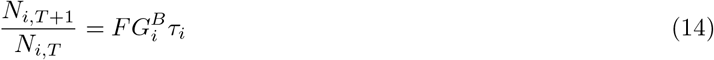

where 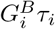 gives the biomass of species *i* at season-end, and *F* converts that biomass to germinated seeds in the next season. Plugging in the expression for *τ_i_* from equation 13 we find

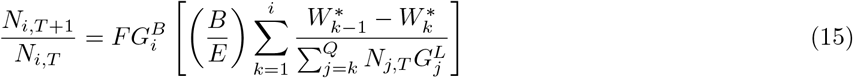

From this equation it is clear that competition is expressed by a reduction in growing season length through consumption of the shared water resource.

#### 3.3 Equilibrium abundances

First note that the sum in the denominator of species 1’s population dynamics expression (equation 15) occurs in each of species 2-Q’s population dynamics equations. This is also true for the sum in species 2’s expression and species 3-Q’s expressions and so on. Therefore by solving for this sum in the expression for species 1, after setting 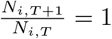 we can iteratively substitute in the solution, eventually solving for 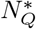.

Setting *N*_1,*T*+1_ = *N*_1,*T*_ in equation 14, we find:

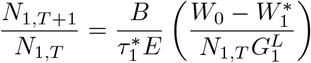

which we solve for the sum in the denominator:

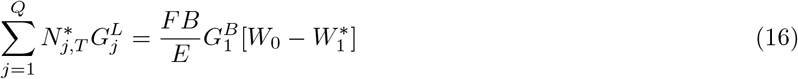

we then substitue equation 16 into our expression for *N*_2,*T*+1_ and solve for 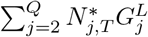, and so on to obtain the following generic expression for 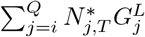:

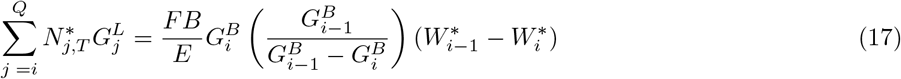

where *G*_0_ = 0. we use this generic expression to solve for the equilibrium abundance of species Q:

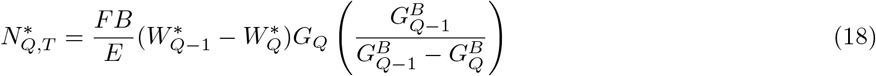

Then plugging this into the expression for species *Q* − 1, and the result into species *Q* − 2, and so on, we find the following expression for the equilibrium population of species *i* ≠ 1, *Q*:

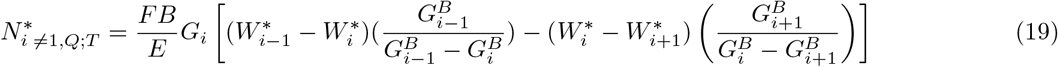

and finally for the first species:

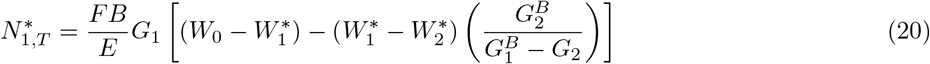

#### 3.4 The unique case of a single-species system

A single species system will snap to its equilibrium population density in a single timestep regardless of its initial density. This is clear when you consider its population dynamic equation:

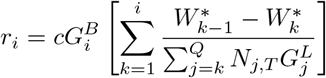

The *N*_1,*T*_’s in the denominators of both sides cancel out giving us:

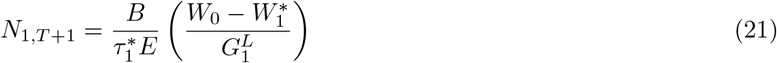

meaning that *N*_1,*T*+1_ is not dependent on the population density of the species in the previous timestep. Moreover, equation 21 gives the equilibrium population density for the species, meaning that at every timestep *T* > 0, species 1 is at equilibrium population density.

#### 3.5 Invasion of a rare type

We consider a species, *q*, growing from low density in a system of *Q* − 1 other species (labelled 1, 2, …, q-1, q+1, …, Q-1) at equilibrium density. By definition, the growth rate of species *q* lies between the growth rates of species *q* − 1 and *q* + 1. Note that following equation 14, the population growth rate of any species, 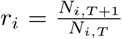, is given by:

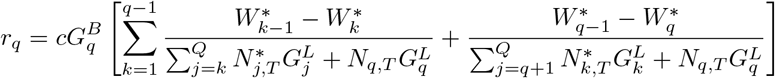

where 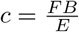. Given that all species *i* ≠ *q* are at population dynamic equilibrium, *r_q_* is given by:

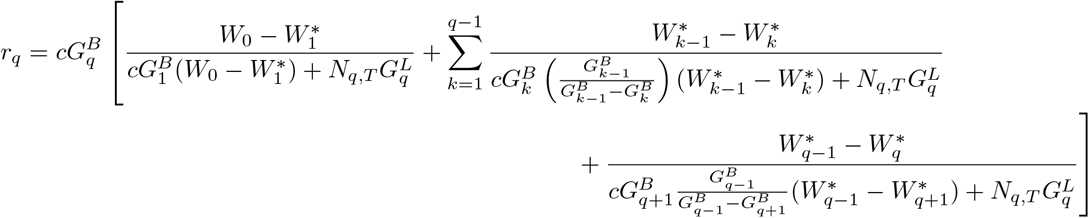

Substituting the expressions from equations 16 and 17 we find:

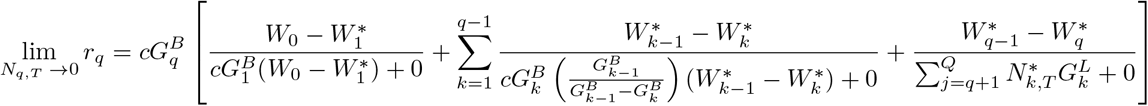

Now, to determine *r_q_* when species *q* is at low abundance, we take lim_*N*_*q,T*→0_*r_q_*_

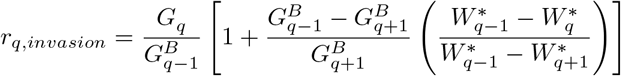

noting that 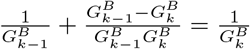, the above expression simplifies to

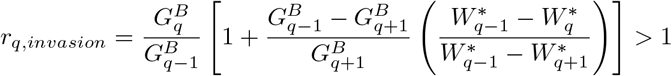

Therefore, invasion of species *q* when rare is governed by

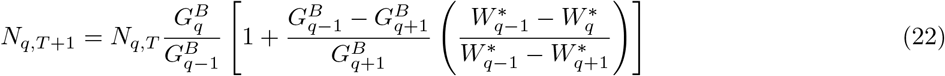

which implies that invasion when rare requires:

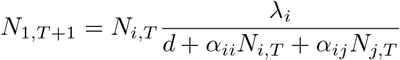

There two special cases, as was true for the equations governing equilibrium abundance: species 1, and species Q. If 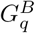 is greater than all other *G^B^*, then its invasion growth rate is given by:

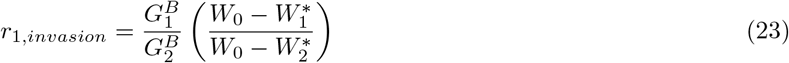

However, if 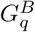 is smaller than other *G^B^*, its invasion growth rate is given by:

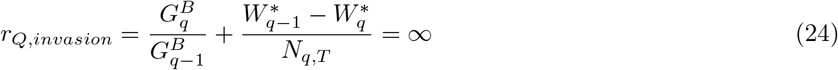

#### 3.6 Decomposition of the per capita competitive effects

Classic phenomenological models of competition among annual plant species partition the effects of competition into intra- and inter-specific interaction terms, labelled *α_ii_* and *a_ij_* respectively. When *a_ij_* and *a_ii_* are large, the Beverton-Holt form of the model, for a two-species system, is written as:

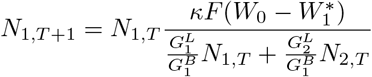

where 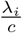 is the growth rate of species *i* in the absence of competition. The population dynamics equation for species 1 can be rewritten to mirror the Beverton-Holt form by dividing the biomass growth rate, 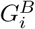 from the numerator and denominator, and setting *d* = 0 to give:

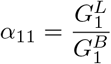

Thereby, in the above:

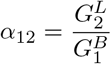

and

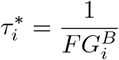

The results are intuitive. Species 1’s intraspecific competitive effect increases as its leaf area growth rate increases relative to its biomass growth rate. Leaf area growth rate is correlated to water consumption, while biomass growth rate is equivalent to a species’ within-season rate of fecundity accumulation. Therefore, the per capita competitive effect of species 1 on itself increases as the ratio of water use to fecundity accumulation increases. Similarly, the interspecific competitive effect of species 2 on species 1 increases as the rate at which it consumes water relative to species 1’s rate of fecundity accumulation increases.

### 4 Using break-even time to understand the invasion condition, positivity condition, and tradeoff curve

We define break-even time, 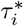 as the growing season length (expressed in conventional time units raised to the B power) which results in a per capita population growth rate of exactly 1 (i.e. 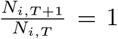). Substituting this condition into equation 14 we find:

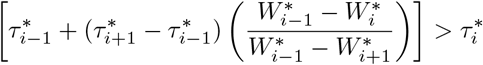

The expression for per capita population growth rate is now rewritten as:

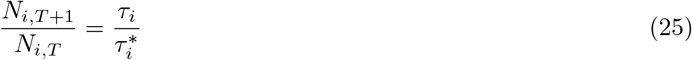

#### 4.1 The invasion condition implies the existence of a positive equilibrium, and vice versa

At equilibrium, 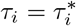 by definition. Now we define the soil water content available to species *i* but not available to species *i* − 1: 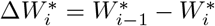 for *i* = 1 through *Q*. We define a similar finite difference for break-even time, except with the indices reversed to avoid considering negative time intervals: 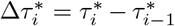. We now rewrite the invasion condition from equation 22:

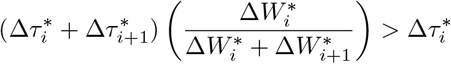

as:

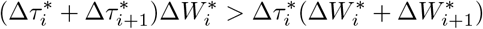

which we rearrange to get:

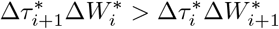

then, multiplying through and subtracting 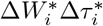 from both sides we get:

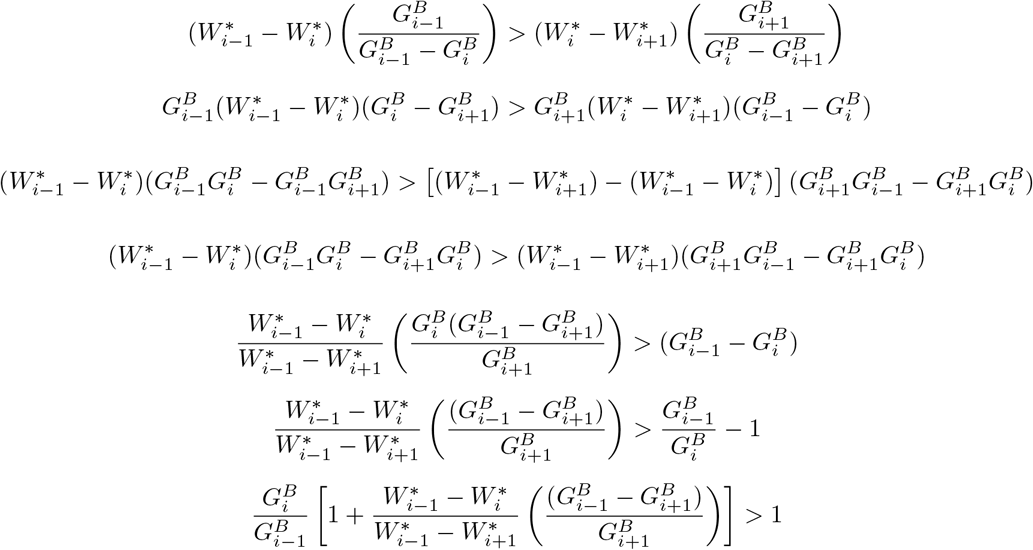

and finally:

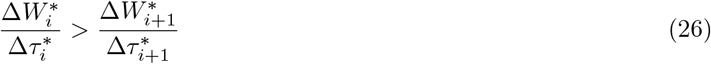

The condition for positive equilibrium density for any species 1 < *i* < *Q* is:

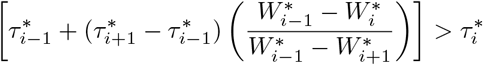

which reduces finally to:

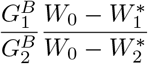

which is identical to the invasion condition for species 1 < *i* < *Q*. Similarly, the invasion condition for the first species is:

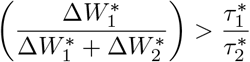

Which in the new notation is:

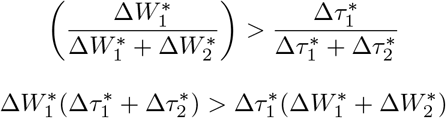

which we rearrange:

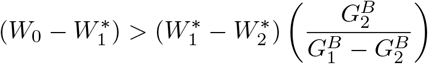

and find, after similar math to the *Q*-species case:

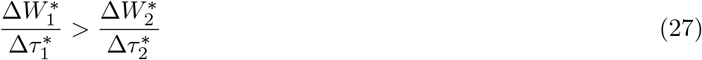

For species 1 to have a positive equilibrium, the following condition must be satisfied (from equation 20):

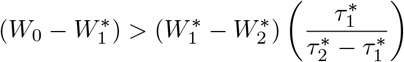

which is equivalent to:

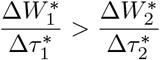

and, using our new difference-based formulation becomes:

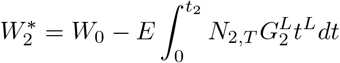

which is identical to the condition in equation 27. Thus, invasion when rare and positivity both require that as we progress through the species from species 1 through Q, each successive jump in break-even time must bring a smaller jump in the amount of water that the species is capable of taking up.

#### 4.2 Break-even time and the growth-phenology tradeoff curve

Now consider a continuous function, *W*(*τ*\*), such that 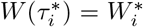 for all possible species. The mutual invasion condition will be satisfied for all successive pairs of species in a system provided *W*(*τ*\*) is both negatively sloped and concave down. We now show that, a tradeoff curve of this form emerges naturally from basic ecophysiological assumptions. We first argue verbally that the physical relationships between soil water potential, volumetric soil water content, and plant hydraulic tension necessitate a negatively-sloped, concave down tradoeff curve. Then we incorporate a stem tissue density – critical water potential relationship into our physiological growth model and show that the resulting tradeoff is of the shape required for coexistence.

We begin with a verbal explanation for why the tradeoff curve between minimum volumetric water content and break-even time must be negatively sloped and concave-down. The tradeoff is expected to have a negative slope because the plant traits that allow transpiration at low levels of soil moisture, such as dense stem tissue that resists embolism [Rosner, 2017], require photosynthate could have been used for productive tissues like leaves, which slows the rate of biomass growth. The positive correlation between 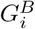 and 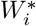, together with the monotonic drying of the soil after the winter rains, implies a tradeoff between biomass growth rate and growing season length, 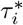, that has a negative in slope.

The tradeoff is also expected to have a concave-up shape primarily because of the nonlinear relationship between soil volumetric water conten, *W* and water potential, *ψ_soil_*. Soil water retention curves relate minus the logarithm of water potential on the vertical axis to volumetric soil water on the horizontal axis. For dry soils, water retention curves have negative slope and are either linear or concave up. A change in units on the vertical axis to minus the soil water potential itself, rather than minus its logarithm, reveals a strongly concave up relationship. Empirical studies imply that a species’ minimum soil potential is a linear function of its stem biomass density [Rosner, 2017]. Now consider three species with equally spaced xylem densities ordered from the lowest density in species 1 to the highest in species 3. The linear relationship between minimum soil water potential and xylem density implies equal spacing of the species along a minimum water potential axis. However, because 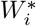 is in units of soil water mass per unit area rather than water potential, and along this axis, the distance between 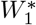 and 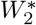 will be smaller than the distance between 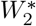 and 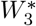.

Because high stem biomass density means slow growth under wet conditions, the break-even times have opposite the order of the 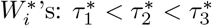. Section 2 also shows that the distance between 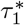 and 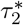 will be smaller than the distance between 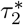 and 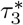 because of the power law allometric growth. Together, the fact that 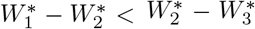 and that 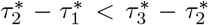 mean that the slope connecting 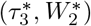 and 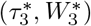 in a plot of soil minimum water threshold versus break-even time, which implies a concave-up tradeoff.

Now, we incorporate a stem tissue density – critical water potential relationship into our growth model. One way that a species can decrease its minimum soil water potential, *ψ*\*, is to invest in thick-walled xylem. In addition to lower critical water potentials, plants which invest in thick-walled xylem have increased wood density as a result, as these structures require more carbon per unit stem volume. The emergent relationship between wood density and minimum soil water content must then be incorporated into our growth model

We incorporate this relationship using equation 8 by assuming that the constant *δ* is composed of two parts, one that converts *L^ν^* to volume, which we call *δ_V_*, and another that converts volume into biomass (stem tissue density), and which is a function of a species’ *ψ*\*, *δ_WD_*(*ψ*\*). As increased wood density is correlated with greater resistance to embolism, *δ_WD_*(*ψ*\*) must be a decreasing function of *ψ*\* (recall that lower values of *ψ*\* mean greater resistance to embolism).We now re-write 8 to reflect this new parameterization, finding:

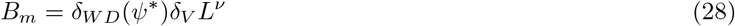

We then substitute the expression for *δ* into equation 9 to find:

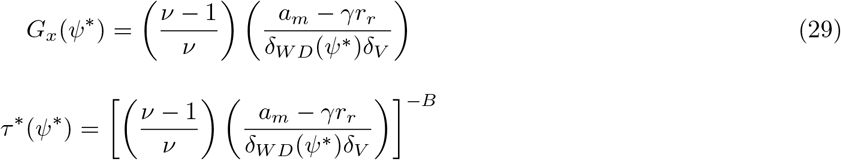

which makes:

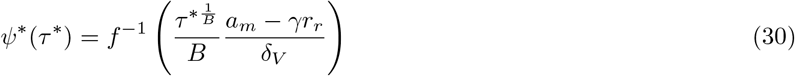

where *f*^−1^(*δ_WD_*(*x*)) = *δ_WD_*(*f*^−1^(*x*)) = *x*. Finally, substituting equation 30 into equation 7 we find:

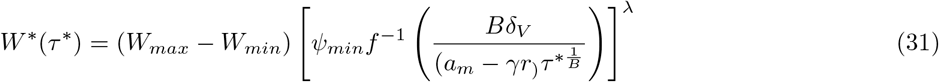

if *δ_WD_*(*ψ*\*) is a decreasing linear function, as empirical work suggests [Rosner, 2017], then equation 31 is a decreasing, concave-up function, as *ψ_min_* < 0 and λ > 0. In fact, as long as *δ_WD_*(*ψ*\*) is a decreasing function, its functional form can vary significantly and equation 31 will remain decreasing and concave-up.

### 5 The graphical method for determining coexistence outcomes

#### 5.1 Invading a single-species population

The water dynamics of any system of competing species are linear in *τ*. More specifically, volumetric soil water content, *W*, is a piecewise linear function of *τ*, where the intervals are defined by species’ season-ends, *τ_i_*. To illustrate, consider a single-species system. Later, we will consider what happens when an early-season species invades, so we label the single species in our system, species 2, to aid notational consistency. Recall that the evapotranspirative dynamics are given by:

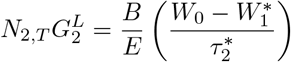

Which, after integrating, becomes:

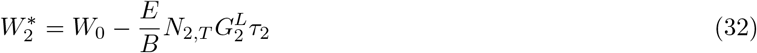

We want an equation for the amount of water availabe at each point in time after *τ* = 0. Because species 2 stops withdrawing water after its season ends at *τ* = *τ*_2_, water content *W*(*τ*) is a piecewise linear function:

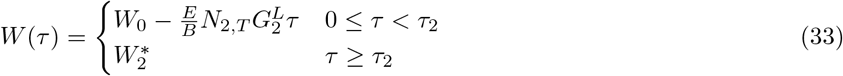

where 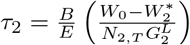. To determine whether a new species can invade this system, however, we want to first determine the equilibrium water dynamics. To do so, we first note that at equilibrium 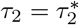. Therefore:

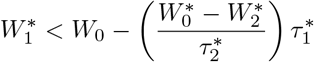

Plugging this into equation 33, we find the expression for *W*(*τ*) when species 2 is at equilibrium:

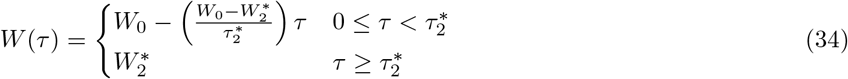

Figure 1B in the manuscript shows the water drawdown trajectory for a single species system at equilibrium. This is equivalent to the resident water dynamics in figure 2 in the manuscript (Figure 2A). Now, consider an invading species, species 1, with a higher minimum water content. This species will be able to invade provided it can grow for at least as long as its break-even time, 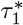, when it is at low density, and species 2 is at equilbrium, as this makes 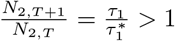. For this to be true, species 1 requires that 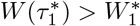, as this means it will still be growing when it reaches its break-even time. Since species 1 has negligible impact on the water dynamics at low density, we need only consider the water consumption trajectory given by equation 34 and shown in figure 1B/2A in the manuscript.

Graphically, we can determine whether 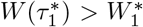 very simply. Figure 1B/2A in the manuscript shows the trajectory of soil water content in the system when species 2 is at equilibrium. To determine 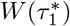, we simply plot a point along the soil water trajectory at 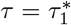. Next we plot a point at 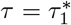 and 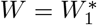. If this point is above the first point we drew, than 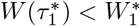, and species 1 will not invade successfully, and the two species will not coexist (Figure 2A in the manuscript). If, however, the second point lies below the first point, then species 1 will successfully invade, and the two species will coexist. Notice that this will be true provided that the second point, 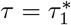 and 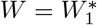, lies anywhere below the line describing the water drawdown trajectory for species 2’s equilibrium.

We can also show this mathematically. For species 1 to invade successfully:

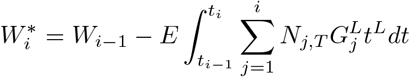

After some algebraic manipulation this becomes:

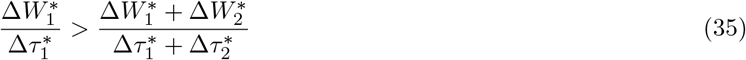

Notice that this inequality reduces to the invasion/positivity condition from section 4.1. The r.h.s of the inequality gives the magnitude of the slope of the first portion of the function in equation 34, or in other words, the rate of water drawdown when species 2 is growing alone and at equilibrium, in *τ* units. This is the slope of the decreasing portion of the soil water content trajectory in figure 2A of the maunuscript. The l.h.s gives the magnitude of the slope of the line connecting the initial soil water water content (*W* = *W*_0_, *τ* = 0) to the end of species 1’s season at equilibrium 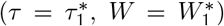 – the same as the second point drawn in determining whether 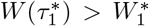 is satisfied graphically, from above. Therefore, the inequality requires that species 2, at monocultural equilibrium, consumes water slow enough that species 1 can reach its break-even time. In other words, the water drawdown rate required to reach species 1’s equilibrium when both species grow together is faster than that required to reach species 2’s equilibrium when growing alone. Note, again, that this is only possible if the point corresponding to species 1’s characteristics 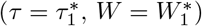 lies below the soil water content trajectory at species 2’s equilibrium.

#### 5.2 Invading a Q-species community

As in the single- and two-species systems, soil water content in a Q-species system is a piecewise linear function of *τ*. With Q-species, the evapotranspirative dynamics are given by:

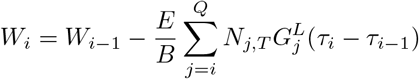

where *t*_0_ = 0. Integrating, we find that:

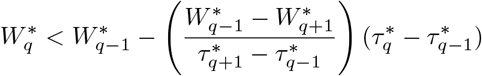

Again, this means that water is consumed at a different during each unique period defined by the number of species still growing. In the first period, all Q species consume water, in the next period when 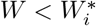 only *Q* − 1 species consume water and so on until species *Q* drops out. The piecewise function defining *W*(*τ*) is:

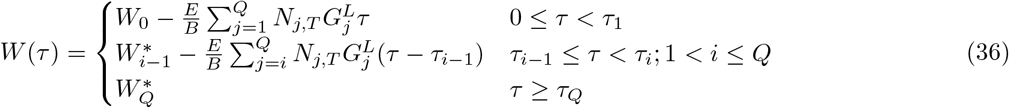

where 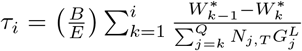. Figure 2B/3A in the manuscript plots an example of these dynamics for a system with 2 species. At equilibrium, soil water content is given by:

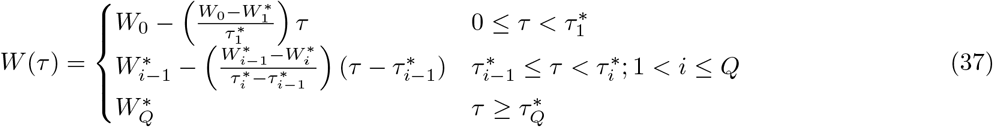

Now consider an invading species with minimum soil water content, 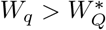. Again, for the invader, species *q*, to increase in population from low density, it must be able to grow for longer than its break-even time. Since species *q* has negligible effect on water consumption at low density, it requires that the water content at its break-even time the soil water content in the resident system under equilibrium exceeds its minimum soil water content threshold, i.e. 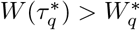.

As in the two-species case, whether 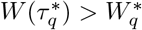 can be determined graphically by first drawing a point along the equilibrium water drawdown trajectory (see Figure 3A in the manuscript) at 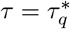, and then drawing a point where 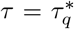 and 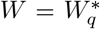. If the second point lies above the first, 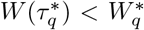, and species *q* will not invade successfully. If the second point lies below the first, 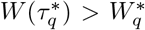 and species *q* will successfully invade. Species *q* will be able to invade provided the point corresponding to its characteristics lies anywhere below the soil water content trajectory for the resident species at equilibrium.

We know show this mathematically by plugging the ineqaulity into equation 37:

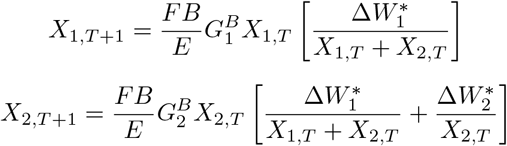

which after rearranging becomes:

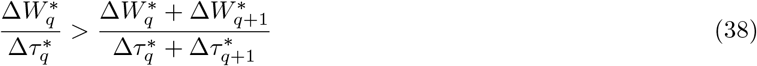

Notice that again, this inequality simplifies to the invasion/positivity condition from section 4.1. The r.h.s is the rate of water drawdown in the period in which species *q* + 1 was the fastest growing species, when the community of all species except *q* are at equilibrium. The l.h.s. is the magnitude of the slope of the soil water content trajectory in the period in which species *q* is the fastest growing species necessary for species *q* to be at equilibrium. Of course, if the l.h.s is greater than the r.h.s. the point which describes species q’s characteristics 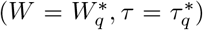 must be below the soil water content trajectory from the resident community at equilibrium.

#### 5.3 An algorithm to determine the coexisting subset of a Q-species system

The inequalities in equations 35 and 38 both simplify to the following expression, first derived in section 4.1:

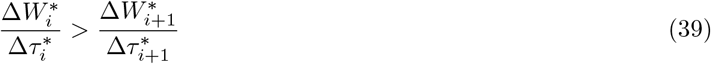

where *τ*_0_ = 0. This inequality must be true for all adjacent species once ordered according to their *τ*s for a community to coexist. If this inequality is not met for all adjacent species, only a subset will coexist. Here we describe a graphical algorithm for determining which subset will coexist using only information about their minimum soil water content and break-even times. In any head-to-head matchup for which the inequality in equation 39 is not satisfied, the species which will persist is the one that consumes water fastest when at monocultural equilibrium, as this species will consume water so fast at densities near-to and greater-than its equilibrium density that its competitor cannot reach its break-even time. In fact, for any set of species for which the inequality is not met for any pair of species, the only persisting species will be the one with the fastest monocultural equilibrium water consumption rate.

To determine the coexisting subset of species from a community for which equation 39 is not satisfied for all adjacent pairs:

1. On a plot with *τ* on the x-axis and *W* on the y-axis, plot points corresponding to the minimum soil water content and break-even times for each species, i.e. points for each 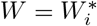 and 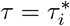, as well as a point at *τ* = 0 and *W* = *W*_0_, which we call the initial point.
2. Draw a line between the initial point and the species characteristic point for which the resulting line has the steepest possible slope. This species will persist.
3. Next, draw a line between that species’ point and the next species’ characteristic point which results in a line with the steepest possible slope.
4. Repeat this process from left to right until no line can be drawn that has a negative slope. All of the species whose characteristic points were connected will coexist.

The result of this process is always the lower convex hull of the cloud of points that includes each species’ characteristic point and the initial point. Each adjacent pair in this set of species will satisfy the requirement in equation 39. It is not the only set of species for which equation 39 is satisfied, but it is the set that will persist because, by maximizing the magnitude of the slope at each step, we guarantee that we select the best competitors for water.

### 6 Time dependent solution for the two species system

To derive the time dependent solution for the population densities in a two-species system we reparameterize the model to reduce its dimensionality. First, we introduce the variable 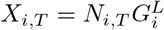 for notational convenience and rewrite equation 14 in the two-species case:

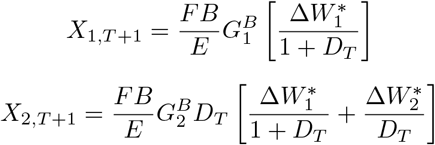

Next, we introduce the variable 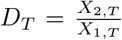 and multiple the left side of the previous expressions by 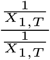 to get:

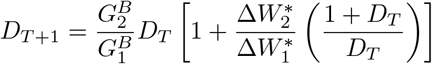

Dividing the expression for *X*_2,*T*+1_ by the expression for *X*_1,*T*+1_ we find that:

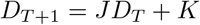

Which, after simplifying becomes:

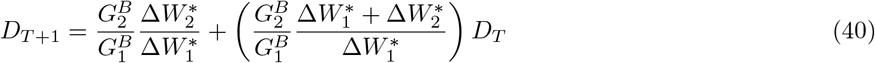

For convenience we set 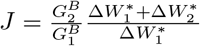 and 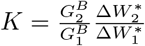 making:

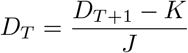

Rearranging slightly:

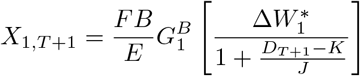

We can now plug this expression back into our expression for *X*_1,*T*+1_ to find:

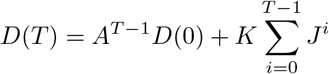

Plugging in 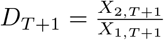 and solving for *X*_1,*T*+1_:

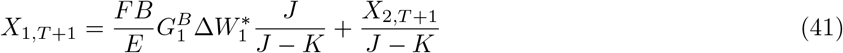

Thus, after the first timestep, the system is constrained to a linear manifold of potential population densities of the two species. This is due to the fact that all of the available water 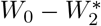 is consumed in each timestep.

Now, since equation 40 is a linear difference equation it can be solved as a function of time for all *T* provided initial conditions are provided.

Using the formula for the solution of a geometric series the above reduces to:

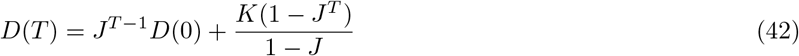

Plugging the above into the expressions for *X*_1,*T*+1_ and *X*_2,*T*+1_ we get the time dependent solutions for the population densities of each species:

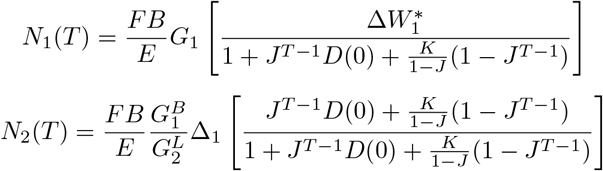

The above can be expressed in terms of more interpretable variables by noting that *K* = *R*_1_ and that

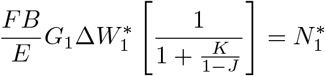

and

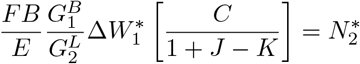

After considerable algebraic manipulation, we find the following expressions for the time dependent population dynamics:

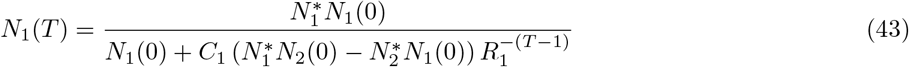

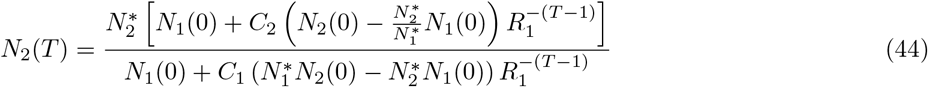

where 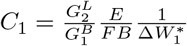 and 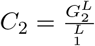

Importantly, the existence of a time-dependent solution for systems with two or fewer species implies that when feasible equilibria exist, they are globally stable.

### 7 Stability analysis

#### 7.1 Deriving the Jacobian

Let’s introduce the parameters 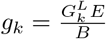, so that we can re-write the dynamics in equation 14 as

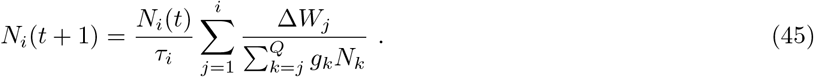

In these new parameters, the formulas for the equilibrium abundances become

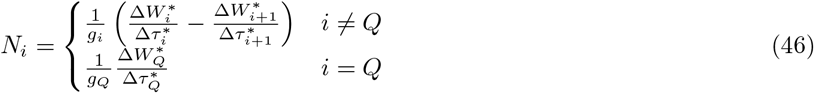

where we are using the parameters 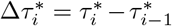 again. From the definitions, we know that 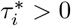 and 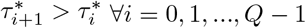 and 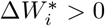. If all of the equilibrium abundances are positive, then we know in addition that 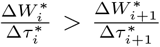. Let’s now compute the Jacobian of the discrete map in equation 45. The derivative of the *i*-th equation by the *j*-th abundance is

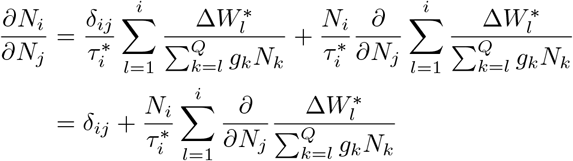

where we used the identity 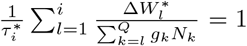 which is true at equilibrium. If *j* < *i* then all the terms in the sum 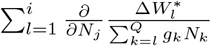 are zero when *j* < *l* ≤ *i*, since there are no *N_j_* in the denominator. Therefore, we compute the derivatives of the terms in the sum only up to the min(*i, j*) term to get that

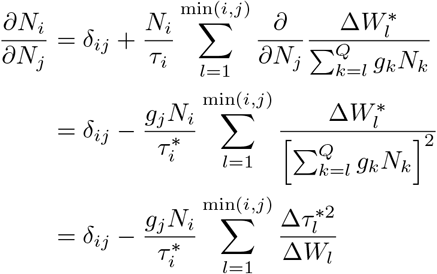

where we used the formula 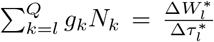 which we derived when computing the equilibrium abundances. Let’s now define 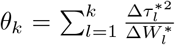 and 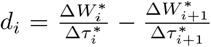 for *i* = 1, …, *Q* − 1 and 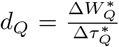. Then, substituting in the abundances from equation 46, we get a formula for the (*i, j*)-th entry of the Jacobian:

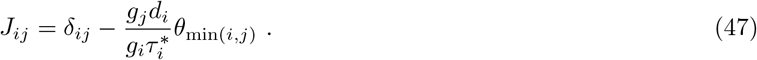

We can also write this formula in matrix form. Let’s define *D* to be the diagonal matrix with *d_i_* as its diagonal entries, *τ_d_* (resp. *g_d_*) to be the diagonal matrix with *τ_i_* (resp. *g_i_*) as its entries and the matrix Θ as

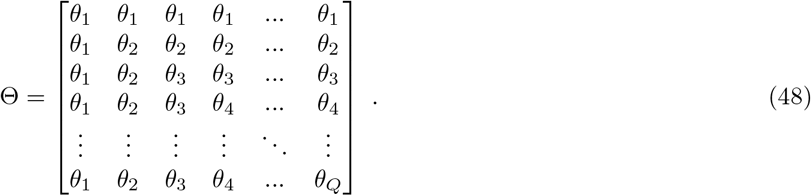

Then, 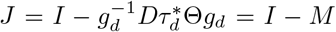 is the Jacobian where we have also defined 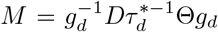. If λ is an eigenvalue of *M*, then −λ + 1 is an eigenvalue of *J*. In the following sections, we will show that the eigenvalues of *M* are contained in the interval (0, 1], which implies that the eigenvalues of *J* are contained in the interval [0, 1) and hence the equilibrium is stable.

#### 7.2 The eigenvalues of *M* are positive and real

First, let’s show that the eigenvalues of *M* are real. The matrix *M* is similar to the matrix 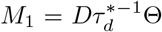 which is in turn similar to the matrix 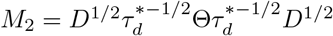. *M*_2_ is a symmetric matrix because Θ is symmetric. Since similar matrices share the same eigenvalues, we know that the eigenvalues of *M* are real because symmetric matrices have real eigenvalues.

The matrix Θ has a special form, which is sometimes called a nested matrix [Stuart, 2015]. If a nested matrix Θ is composed of a strictly increasing sequence of positive real numbers 0 < *θ*_1_ < *θ*_2_ < … < *θ*_*Q*−1_ < *θ_Q_*, then it has a Cholesky factorization Θ = *CC^T^* where *C* = *LK*^1/2^, *L* is a lower triangular matrix with all entries in the lower triangle equal to 1 and *K*^1/2^ is a diagonal matrix with entries 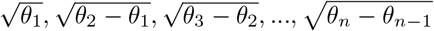 [Stuart, 2015, Horn and Johnson, 2017]. Our *θ*_1_, …, *θ_Q_* sequence satisfies this criterion since it is composed of a cumulative sum of the values 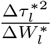. Overall, we find that Θ = *CC^T^* which implies that 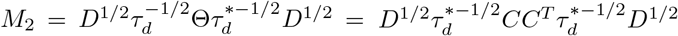. Therefore, *M*_2_ is the product of a matrix with its transpose, since 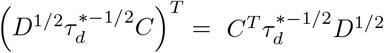, and hence *M*_2_ is positive definite. Since *M*_2_ and *M* share the same eigenvalues, *M* has strictly positive eigenvalues. Therfore, we have shown that the eigenvalues of *M* are on the positive real line.

#### 7.3 The eigenvalues of M are bounded above by 1

*M* (as well as *M_1_* and *M*_2_) are non-negative matrices, which we can see directly from the definitions. Therefore, the Perron-Frobenius theorem applies and all these matrices have an eigenvalue *r* which is the maximum eigenvalue in absolute value ie. *r* > |λ| for all other eigenvalues λ [Horn and Johnson, 2017]. The eigenvalue *r* has a left (resp. right) eigenvector with all positive components, and it is the only left (resp. right) eigenvector with this property, so if we find a left (resp. right) eigenvector of *M*_1_ with positive components, then its corresponding eigenvalue is the Perron root *r* [Horn and Johnson, 2017]. In particular, we will show that 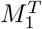 has an eigenvalue of exactly 1 corresponding to a left eigenvector of 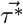.

Let’s check that 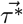 is a left eigenvector of *M*_1_. 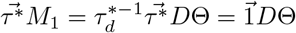 where 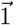 is a vector of all 1’s. So, we want to show that the column sums of the matrix

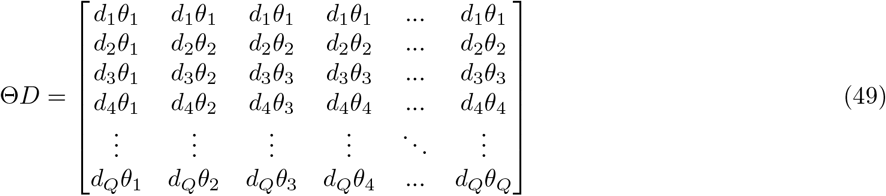

are the same as the entries of the vector 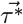. The first column sum is

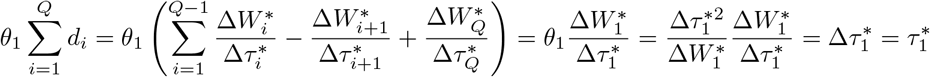

because 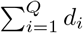 is a telescoping sum. More generally, the *i*-th column sum is

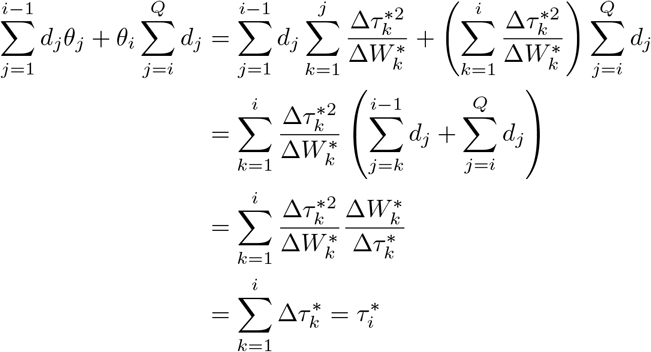

where we used the fact that sums of both the *d_i_* and 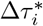 values telescope. Overall, we have shown that the vector 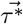 is a left eigenvector of *M*_1_ with eigenvalue 1, which means that 1 is the Perron root of *M*_1_. Therefore, all the other eigenvalues of *M*_1_ are bounded above by 1. *M*_1_ and *M* have the same eigenvalues, so this bound applies to eigenvalues of *M* as well. Since we already know that all of the eigenvalues of *M* are positive and real, we conclude that the eigenvalues of *M* lie in the interval (0, 1], implying that the Jacobian *J* is stable.

### 8 Continuous Equilibrium Population Density in 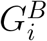

To make notation clearer, we will first define several quantities:

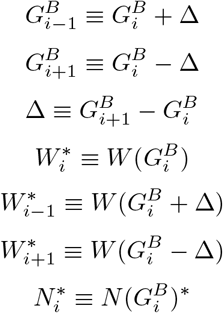

Rewriting equation 14 for species *i* ≠ 1, *Q*:

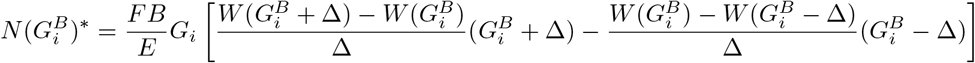

Assuming small Δ, we Taylor expand 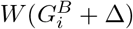 around 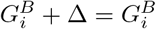, finding:

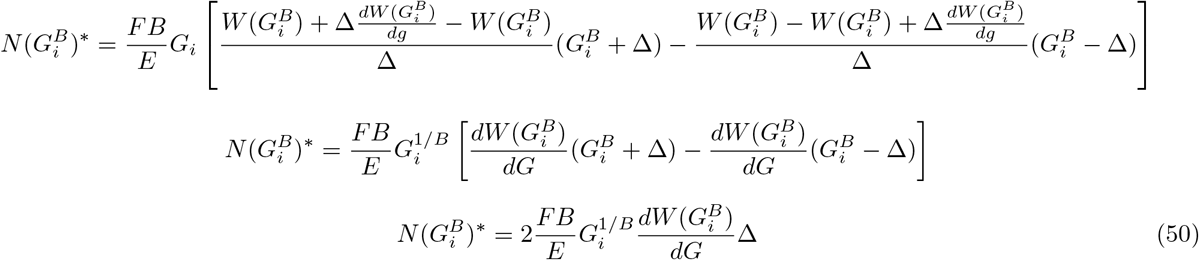

Note that this expression is positive whenever 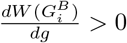 (i.e. the equilibrium abundance of species *i* ≠ 1, *Q* is positive provided that the minimum volumetric water content required for growth is an increasing function of the growth rate).

We now define the equilibrium population density in *G^B^* for species *i* ≠ 1, *Q* as:

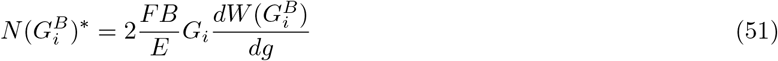

For species 1:

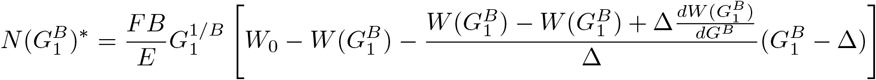

taking lim_Δ→0_:

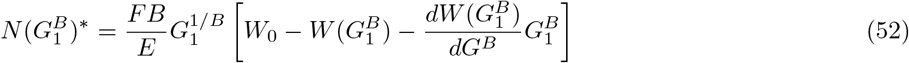

Meaning that 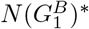 is positive under the following condition:

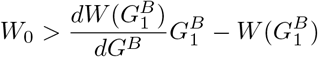

For species Q:

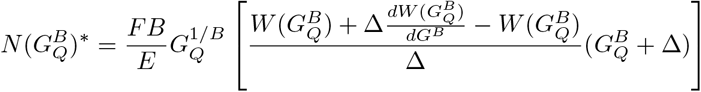

which taking lim_Δ→0_ becomes:

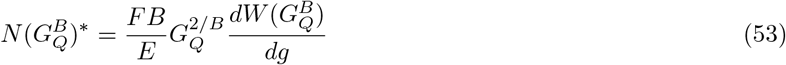

Which is also positive provided 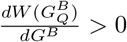

For the case in which 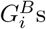 are not evenly spaced we can redefine Δ such that:

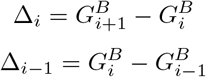

Then, equation 54 becomes:

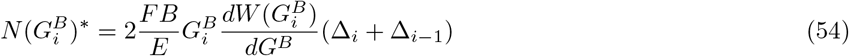

#### 8.1 Probability density function for 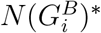

Assume that 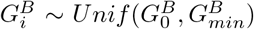 where 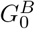 is the *G^B^* corresponding to a species whose minimum volumetric water requirement for growth is *W*_0_ and 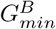 results from a theoretical limit on stem density.

Following from this, we know that:

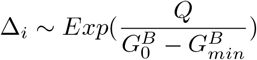

so

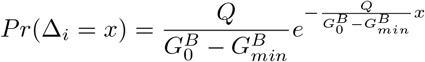

Δ_*i*_ + Δ_*i*−1_ is Erlang distributed (a special case of the gamma distribution) and its probability density function is given by the following:

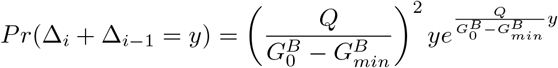

To calculate a probability density function for 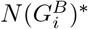, we use the change of variables formula to get:

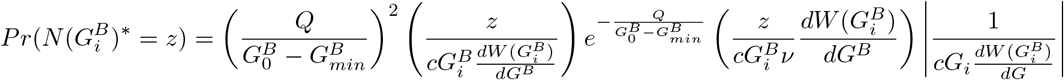

### 9 Model extensions

#### 9.1 2-species model with initial growth period

When species germination occurs before the end of the rainy season,

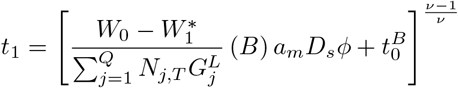

where *t*_0_ is the time at which the rainy season stops. For species 2:

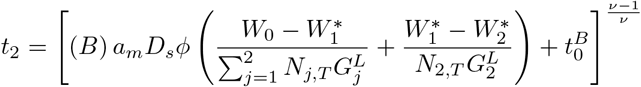

So:

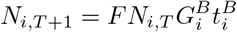

The equilibrium abundances are then:

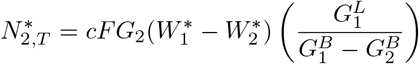

and

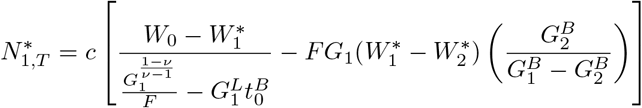

We find the following condition for the coexistence for species 1 and 2:

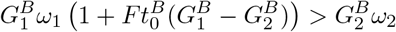

**Table 1:** Parameter Index

| Parameter | Definition | Value for simulations | units |
| --- | --- | --- | --- |
| Physiological parameters |  |  |  |
| $V$ | light availability above the crown | 1.04545 | $kgCm^{-2}yr^{-1}$ |
| $C_a$ | atmospheric carbon concentration | 400 | ppm |
| $C_i$ | internal plant carbon concentration | 275 | ppm |
| $\omega$ | empirical constant in Farquhar photosynthesis model, eq. 1 | 450 | ppm |
| $a_{max}$ | maximum rate of carbon uptake | 0.2 | $kgm^{-2}year^{-1}$ |
| $g$ | plant stomatal conductance | | $kgm^{-2}s^{-1}$ |
| $m$ | empirical constant from Fick's law, eq. 2 | 5.6 | |
| $\Gamma$ | $CO_2$ compensation point | 50 | ppm |
| $\phi$ | shorthand for $\frac{m}{(C_a - \Gamma)(1 + \frac{D_s}{D_0})}$ | | |
| $r_l$ | leaf respiration rate | 0.1 | $kgm^{-2}year^{-1}$ |
| $r_r$ | root respiration rate | 0.05 | $kgm^{-2}year^{-1}$ |
| $\gamma$ | root area to leaf area ratio | 0.5 | |
| Hydraulic parameters |  |  |  |
| $\psi_{leaf}$ | leaf water potential | | $MPa$ |
| $\psi_{soil}$ | soil water potential | | $kPa$ |
| $\psi^*$ | minimum leaf water potential | | $kPa$ |
| $W$ | Volumetric soil water content | | $m^3m^{-3}$ |
| $W_i^*$ | minimum soil water content, corresponds to $\psi^*$ | | $m^3m^{-3}$ |
| $D_s$ | vapor pressure deficit | 1500 | $kPa$ |
| $D_0$ | empirical constant from Fick's law, eq. 2 | 1000 | $kPa$ |
| $k$ | xylem conductivity | 2 | $kgm^{-1}MPa^{-1}year^{-1}$ |
| $E$ | rate of evapotranspiration per unit leaf area | | $m^3m^{-1}day^{-1}$ |
| Allometric and growth parameters |  |  |  |
| $L_a$ | leaf area | | $m^2m^{-2}$ |
| $R_a$ | root area | | $m^2m^{-2}$ |
| $B_m$ | plant biomass | | $kg$ |
| $x$ | arbitrary plant dimension in which growth is linear | | |
| $\delta_{WD}$ | allometric coefficient for relationship between $L$ and $B$ | 0.0815 | |
| $\nu$ | allometric exponent for relationship between $L_a$ and $B$ | 1.5 | |
| $\alpha$ | allometric exponent that relates $x$ to $L$ | 3 | |
| $L$ | allometric exponent relating $x$ to $L_a$, equal to $\frac{1}{\nu-1}$ | | |
| $B$ | allometric exponent relating $x$ to $B_m$, equal to $\frac{\nu}{\nu-1}$ | | |
| $G$ | plant growth rate in $x$ | | |
| $g$ | shorthand for $G^B$ | | |
| Model variables |  |  |  |
| $N_{i,T}$ | population density of species $i$ during season $T$ | | $ind.m^{-2}$ |
| $T$ | year | | years |
| $t$ | time (within season) | | days |
| $\tau$ | time (within season), scaled such that water consumption is linear, $\tau = t^B$ | | |
| $t_i(N_{i,T})$ | The growing season length of species $i$ in standard units, a function of density | | days |
| $\tau_i$ | The growing season length of species $i$ in transformed units | | |
| $\tau_i^*$ | break-even time: the growing season length required for species $i$ to reach L.R.S. = 1 | | |
| $t_0$ | length of initial growth period in model extension | | |
| $F$ | number of germinating offspring produced per unit biomass | | $ind.m^{-2}kg^{-1}$ |
| $N^*$ | equilibrium abundance | | $ind.m^{-2}$ |
| $r_i$ | per capita population growth rate, $\frac{N_{i,T+1}}{N_{i,T}}$ | | |
| $r_{i,inv}$ | low density per capita population growth rate | | |
| $\alpha_{i,j}$ | per capita competitive effect of species $i$ on species $j$ | | |
| $\Delta W_i^*$ | $W_{i-1}^* - W_i^*$ | | |
| $\Delta \tau_i^*$ | $\tau_i - \tau_{i-1}$ | | |
| $X_{i,T}$ | shorthand for $N_{i,T}G_i^L$ | | |
| $D_T$ | shorthand for $\frac{X_{2,T}}{X_{1,T}}$ | | |
| $J$ | shorthand for $\frac{G_2^B}{G_1^B} \frac{\Delta W_1^* + \Delta W_2^*}{\Delta W_1^*}$ | | |
| $K$ | shorthand for $\frac{G_2^B \Delta W_2^*}{G_1^B \Delta W_1^*}$ | | |

**Table 2:** Model summary

| Quantity/Condition | Species 1 | Species $1 < i < Q$ | Species Q |
| --- | --- | --- | --- |
| Break-even time ($\tau_i^*$) | $\frac{1}{FG_1^B}$ | $\frac{1}{FG_i^B}$ | $\frac{1}{FG_Q^B}$ |
| Population growth rate ($\frac{N_{i,T+1}}{N_{i,T}}$) | $\frac{\tau_1}{\tau_1^*}$ | $\frac{\tau_i}{\tau_i^*}$ | $\frac{\tau_Q}{\tau_Q^*}$ |
| Growing season length ($\tau_i$) | $\left(\frac{B}{E}\right) \frac{W_0 - W_1^*}{\sum_{j=1}^Q N_{j,T} G_j^L}$ | $\left(\frac{B}{E}\right) \sum_{k=1}^i \frac{W_{k-1}^* - W_k^*}{\sum_{j=k}^Q N_{j,T} G_j^L}$ | $\left(\frac{B}{E}\right) \sum_{k=1}^Q \frac{W_{k-1}^* - W_k^*}{\sum_{j=k}^Q N_{j,T} G_j^L}$ |
| Equilibrium population density ($N_i^*$) | $\frac{FB}{E} G_1 \left[ (W_0 - W_1^*) - (W_1^* - W_2^*) \left( \frac{G_2^B}{G_1^B - G_2} \right) \right]$ | $\frac{FB}{E} G_i \left[ (W_{i-1}^* - W_i^*) \left( \frac{G_{i-1}^B}{G_{i-1}^B - G_i^B} \right) - (W_i^* - W_{i+1}^*) \left( \frac{G_{i+1}^B}{G_i^B - G_{i+1}^B} \right) \right]$ | $\frac{FB}{E} (W_{Q-1}^* - W_Q^*) G_Q \left( \frac{G_{Q-1}^B}{G_{Q-1}^B - G_Q^B} \right)$ |
| Invasion condition | $\frac{\Delta W_1^*}{\Delta \tau_1^*} > \frac{\Delta W_2^*}{\Delta \tau_2^*}$ | $\frac{\Delta W_i^*}{\Delta \tau_i^*} > \frac{\Delta W_{i+1}^*}{\Delta \tau_{i+1}^*}$ | $W_Q < W_0$ |

